# Soil carbon in the world’s tidal marshes

**DOI:** 10.1101/2024.04.26.590902

**Authors:** Tania L. Maxwell, Mark D. Spalding, Daniel A. Friess, Nicholas J. Murray, Kerrylee Rogers, Andre S. Rovai, Lindsey S. Smart, Lukas Weilguny, Maria Fernanda Adame, Janine B. Adams, Margareth S. Copertino, Grace M. Cott, Micheli Duarte de Paula Costa, James R. Holmquist, Cai J.T. Ladd, Catherine E. Lovelock, Marvin Ludwig, Monica M. Moritsch, Alejandro Navarro, Jacqueline L. Raw, Ana-Carolina Ruiz-Fernández, Oscar Serrano, Craig Smeaton, Marijn Van de Broek, Lisamarie Windham-Myers, Emily Landis, Thomas A. Worthington

## Abstract

Tidal marshes are threatened coastal ecosystems known for their capacity to store large amounts of carbon in their water-logged soils. Accurate quantification and mapping of global tidal marshes soil organic carbon (SOC) stocks is of considerable value to conservation efforts. Here, we used training data from 3,710 unique locations, landscape-level environmental drivers and a newly developed global tidal marsh extent map to produce the first global, spatially-explicit map of SOC storage in tidal marshes at 30 m resolution. We estimate the total global SOC stock to 1 m to be 1.44 Pg C, with a third of this value stored in the United States of America. On average, SOC in tidal marshes’ 0-30 and 30-100 cm soil layers are estimated at 83.1 Mg C ha^-1^ (average predicted error 44.8 Mg C ha^-1^) and 185.3 Mg C ha^-1^ (average predicted error 105.7 Mg C ha^-1^), respectively. Our spatially-explicit model is able to capture 59% of the variability in SOC density, with elevation being the strongest driver aside from soil depth. Our study reveals regions with high prediction uncertainty and therefore highlights the need for more targeted sampling to fully capture SOC spatial variability.

## Main

Tidal marshes, like other blue carbon^1,2^ ecosystems (mangroves and seagrasses), are a global soil organic carbon hotspot owing to high rates of autochthonous and allochthonous organic matter deposition and slow decomposition in waterlogged soils. In addition to securing this carbon over millennia, tidal marshes are also considered to be one of the most effective ecosystems for carbon accumulation^3^. Their capacity to grow volumetrically with sea level rise means that, in comparison to terrestrial ecosystems, carbon saturation is far less likely to occur, providing continuous climate change mitigation benefits.

Tidal marshes were estimated to cover an area of 52,880 km^2^ in 2020, distributed across 120 countries and territories^4^, however this is only a fraction of prior extent. It is likely that over 50% of global tidal marsh habitat has been lost since 1800^5^, with modern annual loss rates of between 0.2 – 2%^6–8^ resulting from global warming, sea-level rise, and anthropogenic activities such as agricultural or urban expansion, and human engineering of coastal floodplains and river systems^9^. Owing to the many ecosystem services tidal marshes provide, as well as their role in climate change mitigation, there is an increasing need to conserve and restore tidal marsh habitats globally^10,11^.

As part of effectively managing tidal marshes and quantifying their climate mitigation potential, there is a need to first understand the quantities of carbon stored within their soils^12^. Though we have information on SOC in tidal marshes at local to regional scales in areas like the conterminous United States^13^, Great Britain^14^ and Australia^15^, we currently lack a global scale analysis supported by extensive field-based observations and scaled up beyond temperate marshes^16^. Without this information, the scientific community and practitioners have to rely on global averages that are not ecosystem-specific^17^ and that are highly biassed toward temperate regions where data is more abundant^18^, or they must collect resource-intensive in-situ field measurements.

Here, we present the first global spatial model of SOC stock in tidal marshes and sampling bias associated uncertainties. We did this by coupling a higher tier global tidal marsh extent map^4^ with a new global dataset containing 3,710 measurements of tidal marsh soil properties and carbon content^19,20^, and using a machine-learning approach including environmental covariates identified by expert elicitation as potential drivers of soil carbon density. To align with IPCC recommendations, we present spatially-explicit SOC stocks to 1 m depths. This depth profile is used to represent soil more susceptible to emissions linked to disturbances. We also provide estimates for both the 0-30 cm and 30-100 cm soil layers, to align with both the numerous SOC studies that are confined to the upper layers and to match the conventional 30 cm depth for terrestrial carbon estimates.

## Global distribution of tidal marsh soil organic carbon

Our spatially-explicit model predicts 1.44 Pg C in the top metre of tidal marsh soils globally. It likely represents a considerably more spatially accurate estimate than previous efforts, given that the model is trained on a globally distributed empirical dataset^19,20^ and hypothesis-driven landscape-level drivers (Table S1). Previous global tidal marsh carbon stock estimates have taken a wide range of values. With lower values such as the 0.43 ± 0.03 Pg C estimated in the top 0.5 m from a dataset based mostly on North American tidal marshes^18^, continental SOC averages to 1 m multiplied by extent estimates (1.41–2.44 Pg)^21^, or ranging between 0.86 and 1.35 Pg C^16^ estimated to a depth of 1 m from the SoilGrids map, a global machine-learning map from agricultural soils and terrestrial ecosystems data^17^. Conversely, simple calculations based on an average SOC value, applied to an overestimated tidal marsh extent have indicated that the global stock could be as high as 6.5 Pg C^22^. Our improved prediction of total global SOC in tidal marshes is significantly lower than this upper estimate, with our model predicting a range of 0.87-1.62 Pg C. It should be noted that our global tidal marshes SOC stock estimate is conservative because the underlying global map of tidal marshes we used only extends to 60°N and there are considerable unmapped areas of tidal marsh in the Arctic^4^.

The magnitude of regional and national carbon stocks is strongly driven by tidal marsh area. For example, a high proportion of the total global marsh carbon was located in the Temperate Northern Atlantic (Fig. 1a), which holds almost half (45%) of the global tidal marsh extent^4^. Countries with the highest predicted total SOC in tidal marshes (the U.S., Canada, and Russia, followed by Argentina, Australia, and Mexico – Fig. 1b, Table S2) had both high carbon per unit area and large marsh extents^4^. Our country-level tidal marsh SOC stock estimates are in line with those found in several regional studies. For example, stocks of 720 Tg C to 1 m were projected in the conterminous United States^13^, which is comparable with our value of 520 Tg C. Similarly, 5.2 Tg C was estimated for the shallow (28 ± 16 cm^23^) tidal marsh soils of Great Britain^23^, which aligned with our prediction for the top 30 cm being 4.7 Tg C. This highlights the importance of geographical context when considering our findings. For example, many marshes across Great Britain have relatively shallow soil profiles^23^, and as such our estimate to 1 m would overestimate the country stock. Conversely, in settings with a longer history of relative sea-level rise or very high tidal ranges, marsh soil can reach depths exceeding 1 m^24,25^, and thus we may underestimate total SOC stock.

**Fig. 1.**
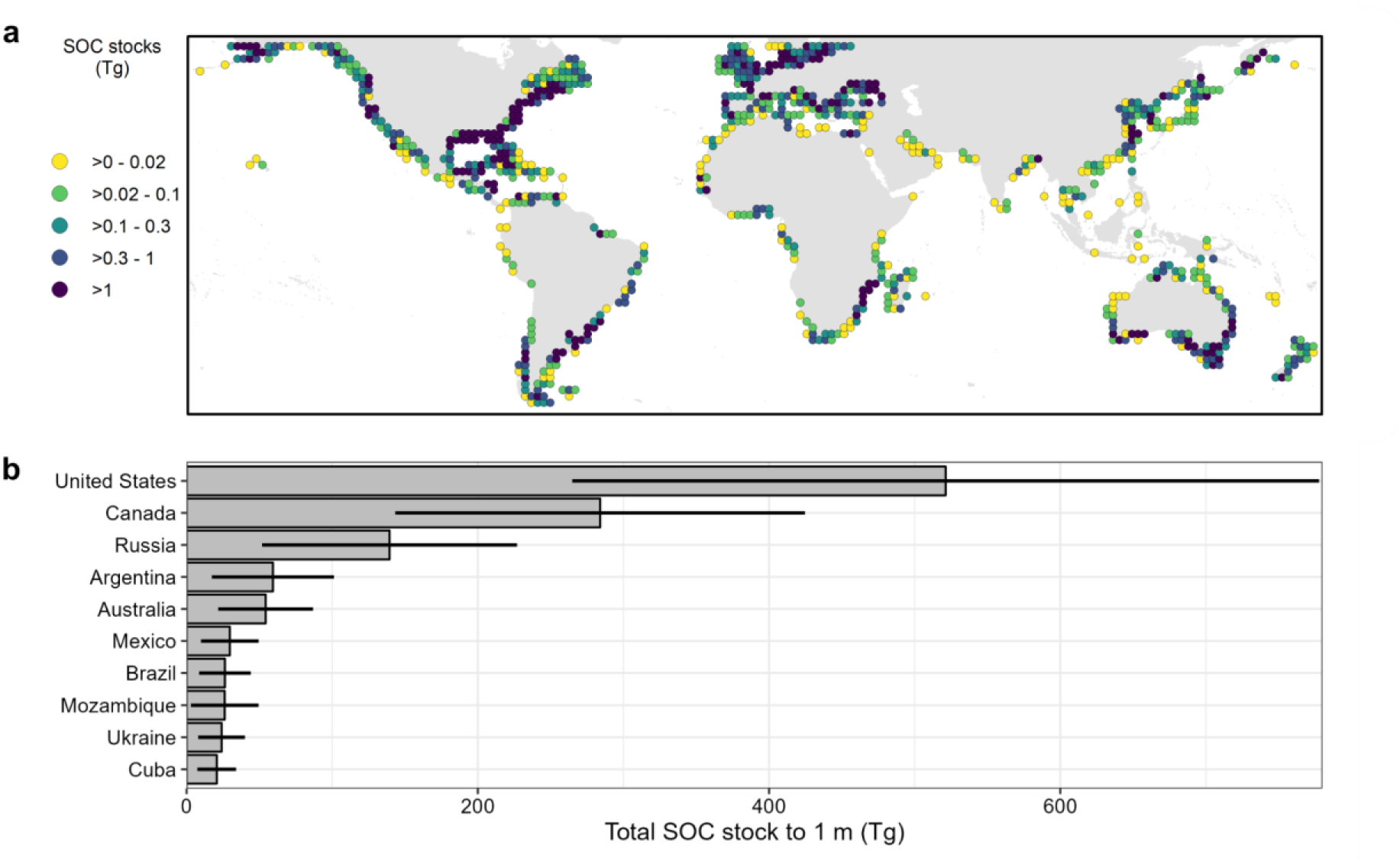
| Total soil organic carbon stocks (SOC, Teragrams) in the top 1 m of tidal marshes. a) aggregated per 2° cell, and b) for the ten countries with the highest total SOC stock. Values refer to predicted SOC stocks after removing pixels outside the area of applicability (AOA), i.e. where we enabled the model to learn about the relationship between SOC stocks and the environmental drivers. Whiskers represent expected model error.

We predicted that the average SOC per hectare in tidal marshes globally is approximately 83.1 Mg C ha^-1^ in the 0-30 cm layer and of 185.3 Mg C ha^-1^ in the 30-100 cm layer (Fig. 2), with an average predicted error of 44.8 Mg C ha^-1^ (Fig. S3) and 105.7 Mg C ha^-1^ (Fig. S4), respectively. Our value to 1 m of 268 Mg C ha^-1^ refines previous global estimates, such as 162 Mg C ha^-1^ derived from local carbon data of unclear origin^22^, and 317 Mg C ha^-1^ averaged from an undefined number of studies^26^. Our approach accounts for the spatial variability in carbon and is an improvement upon averages based on reported values alone. Further, our central estimate for tidal marsh soils is within the range of those predicted for mangrove soils (232-470 Mg C ha^−1^)^27^, confirming that these blue carbon ecosystems store significantly more SOC per unit area than many terrestrial ecosystems^28^.

**Fig. 2.**
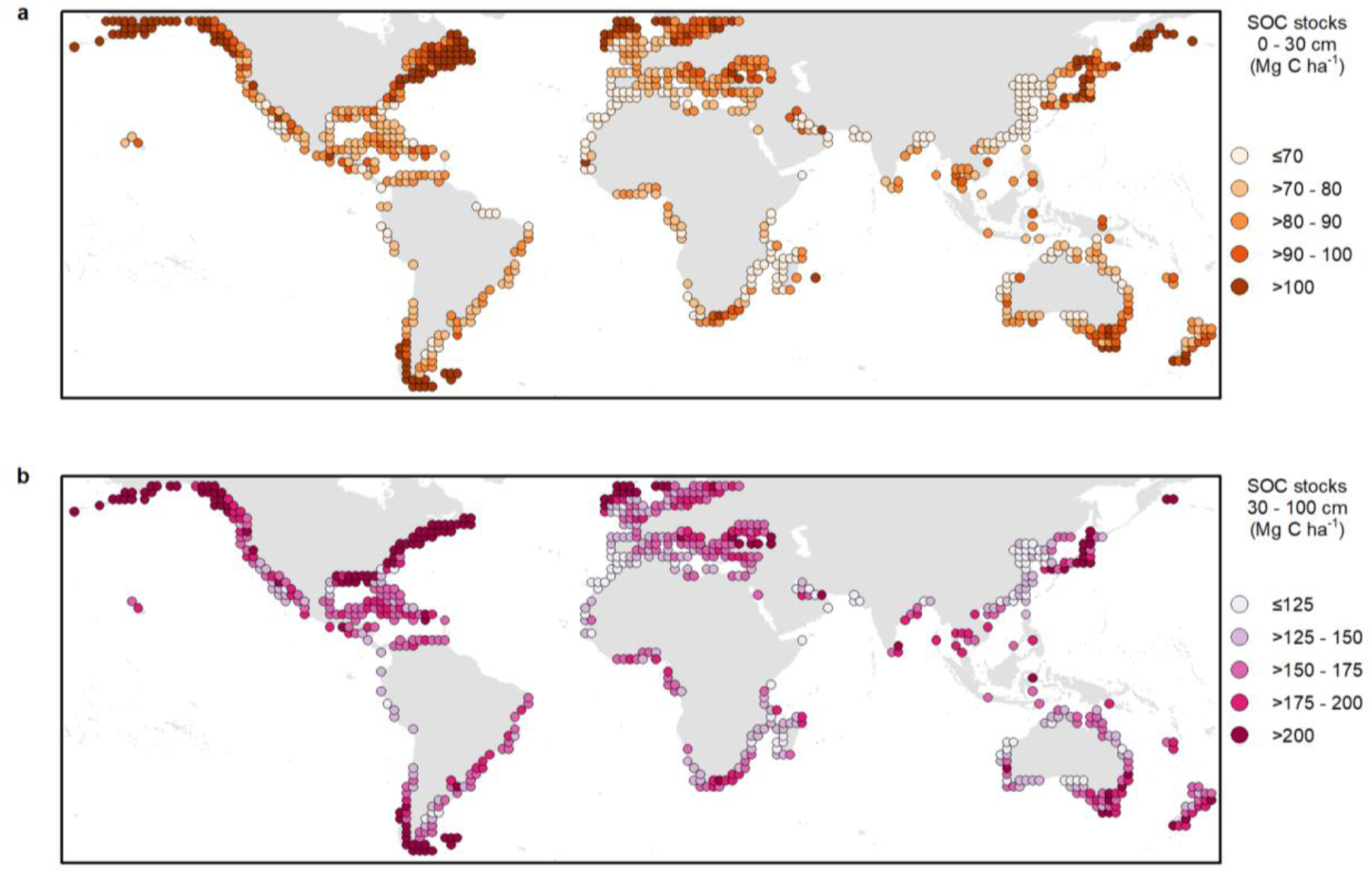
| Global distribution of tidal marsh soil organic carbon (SOC) for a) the 0-30 cm soil layer and b) the 30-100 cm soil layer (aggregated per 2° cell). Values refer to predicted SOC per unit area (Mg C ha^-1^) after removing pixels outside the area of applicability (AOA), i.e. where we enabled the model to learn about the relationship between SOC and the environmental drivers. Cells with 0 % of pixels within the AOA are not displayed. Because fewer training data points were available in the deeper soil layer, more pixels are outside the AOA and thus fewer cells are displayed in the lower panel. Initial predicted values and the proportion of pixels in each cell within the AOA are presented in Fig. S1 and Fig. S2.

Overall, our analysis indicates larger SOC per hectare in higher latitudes, with northern areas of the Temperate Northern Atlantic and Pacific realms having particularly high values (Fig. 2). At the regional level, within both depth layers, temperate areas generally have slightly higher average SOC values than tropical regions, although there is high expected model error associated with each prediction (Fig. S3, Fig. S4). This finding goes against the hypothesis that higher temperatures are generally associated with higher SOC^29^, due to the increase of productivity and growth of vegetation^30^. Instead, the lower soil temperature could limit SOC breakdown enhancing its storage potential^30^, or more generally temperature could be a weak driver at the global scale^31^. The high tidal marsh SOC predicted to 1 m in higher latitudes is somewhat impacted by limited training data and a low proportion of our predictions in the area of applicability (Fig. 3). Although not included as drivers in the model, the larger SOC in higher latitudes could be explained by an increase in vertical land movement in these areas, which leads to the decrease in the relative sea-level rise^32^, impacting the lateral distribution of tidal marshes and their capacity to slow decomposition and ability to accumulate carbon.

**Fig. 3.**
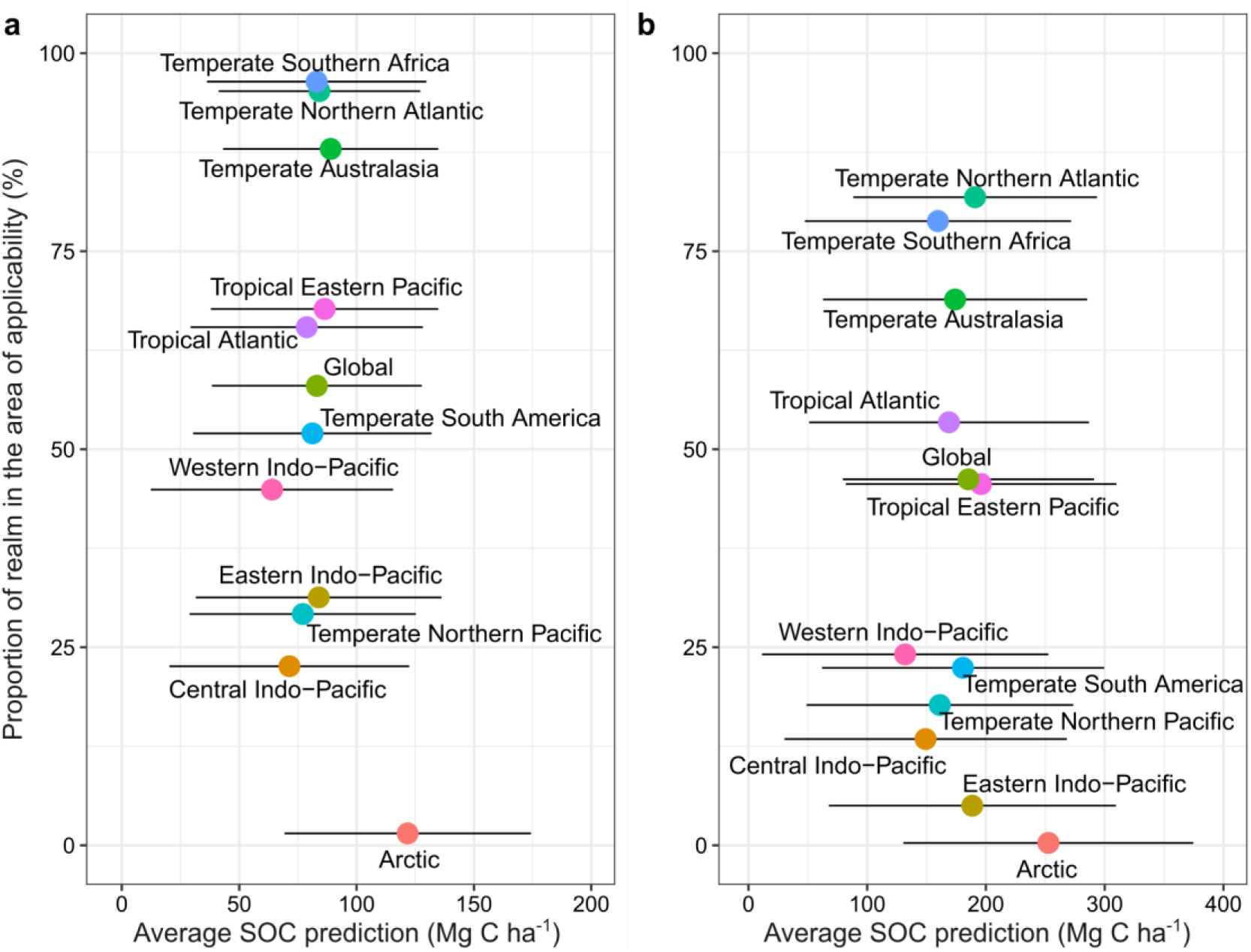
| Realm level summary statistics of soil organic carbon (SOC) in a) the 0-30 cm soil layer and b) the 30-100 cm soil layer. For each soil layer (0-30 cm and 30-100 cm), the x-axis shows the proportion of the realm within the area of applicability (AOA), i.e. where we enabled the model to learn about the relationship between SOC and the environmental drivers, and the y-axis shows the average final predicted SOC, after masking out areas outside the AOA. Whiskers represent the expected model error for each prediction. Colours are mapped to realms, which correspond to the biogeographical realms of the Marine Ecoregions of the World^33^, and the global average.

## Drivers of soil organic carbon in tidal marshes

Overall, our model performed well in describing variation in our SOC training data, with an R^2^ of 0.59. We selected hypothesis-driven environmental drivers (Table S1) to limit the number of covariates included, as well as a spatial cross-validation strategy to avoid issues of overfitting^34^. Soil depth was the most important driver of SOC density (Fig. 4). This is consistent with findings quantifying SOC in mangrove ecosystems^27,35^. Depth has a strong influence on SOC concentration and bulk density, the interaction of which results in relatively stable SOC density measurements across the soil profile^13,19^. Elevation was also an important variable, with higher SOC values predicted at lower elevations, which are generally characterised by higher sedimentation rates allowing more trapping of organic C^36^, as well as more frequent inundation providing opportunity for deposition of tidally-distributed sediments to settle on tidal marsh surfaces^37^ and for plants and their roots to grow better, adding organic material to the soils. Maximum SOC storage occurs above mean sea level where i) tidal marsh vegetation can thrive and serve as a C source, ii) vertical space (or accommodation space) remains available for accumulation of organic matter enriched sediments from autochthonous and allochthonous sources, and iii) decomposition of SOC is hampered by the anaerobic conditions created by higher inundation frequencies^31^. While the variable importance of the model gives an indication of the variables driving the patterns of SOC, it should be interpreted cautiously as underlying interactions between variables are not captured.

**Fig. 4.**
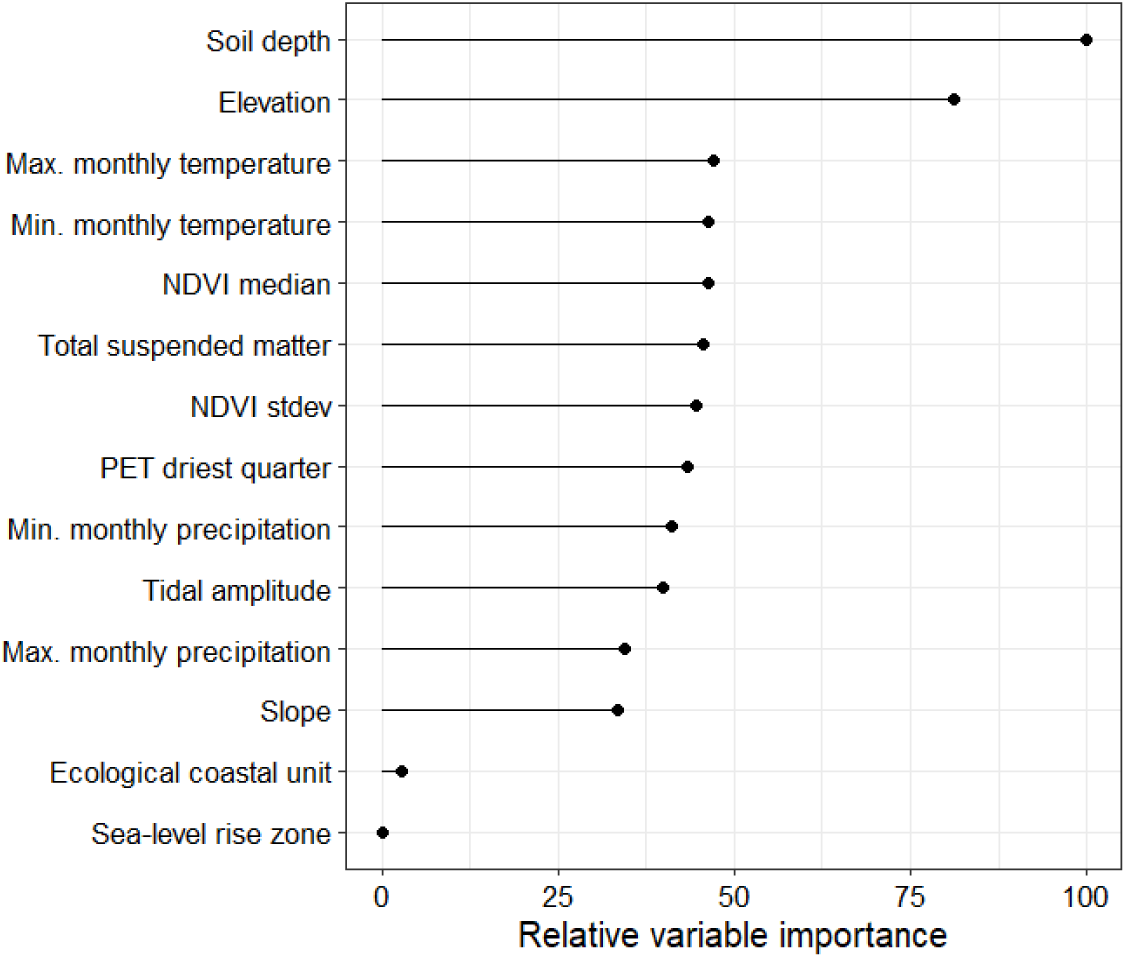
| Variable importance of the random forest model used to make the predictions of tidal marsh soil organic carbon stocks. NDVI, “Normalised Difference Vegetation Index”, PET, “potential evapotranspiration”. Importance was set to “impurity” in the model settings.

Our model includes the best globally available environmental covariates used to predict SOC in tidal marshes. However, there are a number of broad-scale drivers identified as potentially important predictors of soil carbon density which were poorly represented in our model or for which there were no globally available data products. For example, sea-level rise history has been linked to carbon storage in marshes^31,38^. This relationship was not apparent using our modelling method, but that is likely driven by the relatively coarse-scale classification of sea-level rise history zones^39^ used in the analysis. While we included the Normalised Difference Vegetation Index (NDVI) as a proxy for tidal marsh vegetation type and the source of SOC, data sources representing the distribution of dominant plant species, diversity, or species assemblages at a global scale would be an important covariate to develop^40^, as carbon stocks can vary with species^41,42^. Additionally, a high resolution map of coastal typology of tidal marshes would help refine predictions as geomorphic setting (for example deltas, estuaries, lagoons, composite deltas and lagoons, as well as sediment conditions and geology) controls the SOC stock via the type and rate of sediment supply to the coastline, nutrient loading/limitation, and organic matter diagenesis^43^. A global map of shoreline morphology would also better represent the accommodation space available for carbon storage^31,44^, rather than using coastal typology.

## Locations for priority sampling

While our global tidal marsh SOC model is underpinned by an extensive training dataset^19,20^ of 42,741 observations from 3,710 unique locations (Fig. S5), the applicability of the model output is reduced in some regions where data is scarce or inexistent. Over 85% of the training locations are from the U.S.A. (n = 2,005), the U.K. (n = 944) or Australia (n = 284). By implementing an area of applicability (AOA) method^45^ we restrict predictions to locations where the relationship between the training data and environmental covariates is meaningful. Due to the sparsity of training data from some locations, areas where our model predictions are robust (the AOA) represent 58% of mapped marshes globally for the 0-30 cm layer and 46.2% for the 30-100 cm layer.

The spatial applicability of our model varies between regions, with high AOAs (>85%) for many temperate regions, but very low applicability for the Arctic and much of the tropics (Fig. 3, Table S3). By understanding this interplay between the predictions and their uncertainty, our analysis identifies priority areas for sampling to better parameterise future models (Fig. 5). For example, our analysis predicts high SOC across the high Arctic; however, this region is characterised by limited training data and thus high per pixel expected error. In our final analysis these areas are outside the AOA, and therefore we have significant uncertainties when estimating SOC in this region. Many temperate areas were also predicted to have high SOC, but here predictions were underpinned by extensive field measurements and thus subject to greater certainty. Finally, many tropical and subtropical areas of tidal marsh are predicted to have low SOC. Tidal marshes in these regions, where mangroves are more likely to occur, are critically understudied^46^, and our predictions in these regions are limited by our knowledge on their structure and extent. SOC in tropical tidal marshes has been shown to be very variable^47^ and as such our model predictions will be sensitive to the limited training data available; however, the currently available studies suggest generally lower SOC per unit area than the global average^48^.

**Fig. 5.**
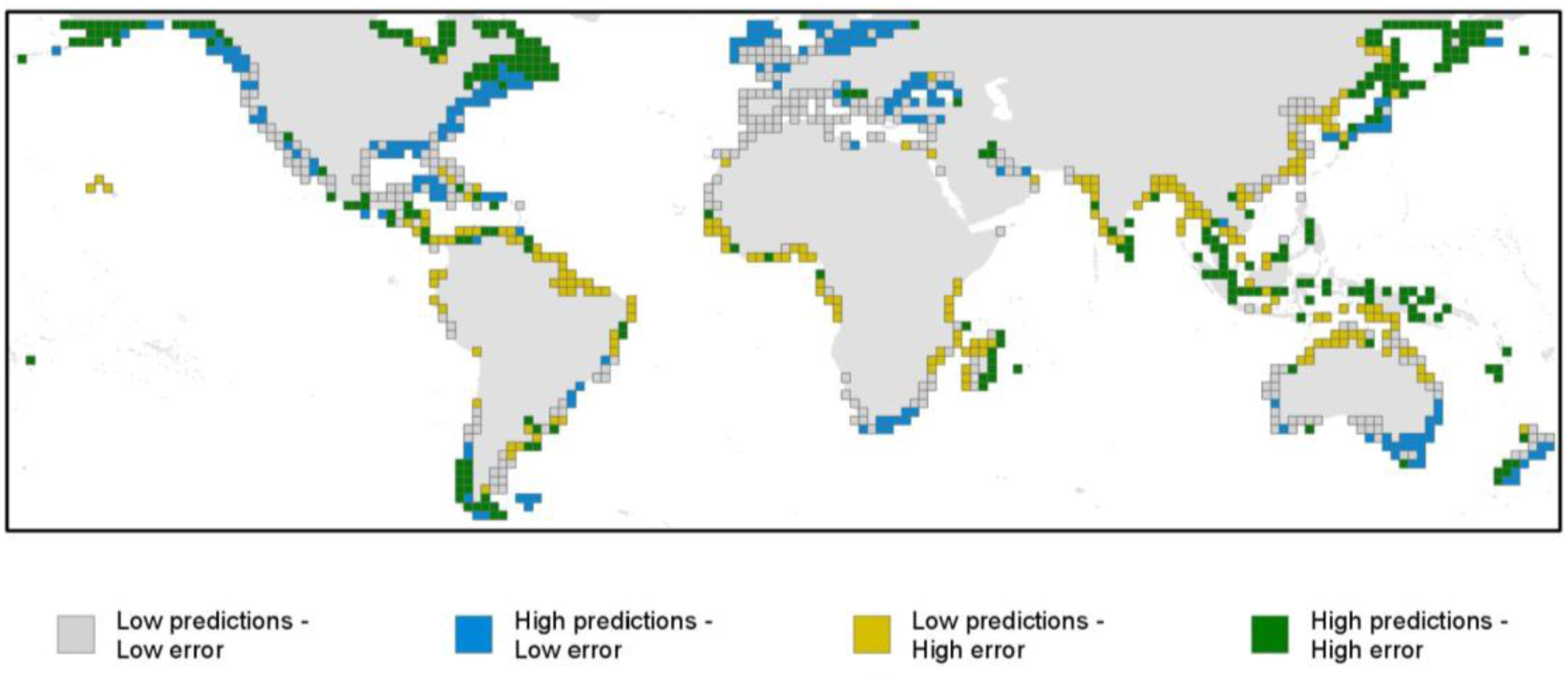
| Bivariate plot showing predicted SOC stocks per unit area and expected error. Values are for the initial model predictions and expected model error (i.e., not masked by the area of applicability), aggregated per 2° cell. The plot shows locations with low predictions and low error (grey), high prediction and low error (blue), low predictions and high error (yellow), and high predictions with high error (green).

Our research highlights two key pathways for future work, firstly greater SOC field measurements from the extensive areas of Arctic tidal marsh to better quantify potential critical stocks in those regions, areas which are significantly threatened by climate change^49^. Secondly, we require far greater understanding of tidal marshes within the marsh-mangrove ecotone in the tropics to elevate their role as a key blue carbon ecosystem.

The uncertainties in our model also have a depth component, with consistently reduced AOAs in the 30-100 cm layer compared to the 0-30 cm layer (Fig. 3). This reduction is driven by the smaller number of data points in this deeper layer (35 % of all data). This depth disparity in data coverage could be addressed by more consistent sampling to 1 m across studies; however, it is also important to note that not all marshes have an organic matter layer that exceeds 30 cm depth, such as in the northern Pacific coast^50^ and Great Britain^23^. These shallow marshes formed more recently atop tidal flats – thus the deeper layers do not hold levels of SOC similar to deeper layers of older marsh soils soil layers. This has implications when estimating SOC stocks to 1 m following IPCC guidelines, and as such we advocate providing estimates across both the 0-30 cm and 30-100 cm layers.

## Outlooks and policy implications

Tidal marshes hold a substantial stock of organic carbon, and our globally consistent map improves knowledge of coastal ecosystem services while practically supporting the inclusion of tidal marsh blue carbon in national inventories that could be used as a baseline for carbon accounting. Such data could catalyse coastal ecosystem conservation and restoration efforts. Disentangling sources of error from the available training data allows a greater understanding of prediction uncertainty, while our spatially explicit analyses of expected model error clearly highlights locations where future work is urgently needed. These areas include the Arctic and the tropics, and this work should include targeted field data collection of tidal marsh extent, structure and carbon stocks. Our dataset further enables better quantification of potential losses in SOC from land use change or conversely benefits from conservation and restoration. International interest in blue carbon ecosystems is high, strongly driven by the calls for climate action under the United Nations Framework Convention on Climate Change (UNFCCC) and the UN Sustainable Development Goals. The very high carbon content and high sequestration rates of tidal marshes places them at the centre of growing efforts to mitigate climate change through avoiding future losses, securing current stocks and also through ecosystem restoration^11,21^. A growing number of countries are including such ecosystems in their national commitments to climate change mitigation as part of their Nationally Determined Contributions, where our study can fill in significant gaps where data is scarce or inexistent.

## Methods

### Tidal marsh soil carbon training data

The global database was compiled from two main sources: the recently produced global tidal Marsh Soil Organic Carbon (MarSOC) Dataset^19,51^ (https://zenodo.org/records/8414110) and from the Coastal Carbon Research Coordination network^20,52^ (https://ccrcn.shinyapps.io/CoastalCarbonAtlas/). The data was filtered to those sites located between 60°N to 60°S due to the environmental covariate data being limited to this region, and within a coastal zone data mask^53^. This data contains 42,741 points from 22 countries and 3,710 unique locations^54–256^.

### Spatial modelling of soil organic carbon

We used a 3D approach to model organic carbon density (OCD) to maximise the applicability of the collected data and reduce the need to make assumptions about OCD trends along the soil profile^257^. We thus modelled OCD as a function of depth (*d*), and a series of environmental covariate layers (*X_n_*):

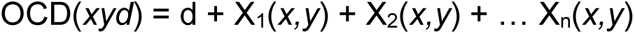

where *xyd* are the 3D coordinates in decimal degrees of latitude and longitude, and soil depth (measured at the centre of a sampled soil layer). The resulting spatial prediction model can then be used to predict OCD at standard depths of 0, 30, and 100 cm, so that the soil organic carbon stock (SOC) for the 0-30 cm and 30-100 cm soil layer can be calculated using their respective layer thicknesses^257^:

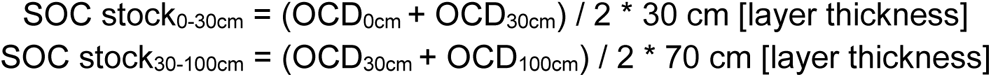

#### Environmental covariates

Environmental covariates were used based on hypothesised landscape-level drivers of soil organic carbon in tidal marshes (Table S1). Drivers were selected by the authors whose expertise includes regional and field-based knowledge as well as considerable prior knowledge of global-scale modelling. This was an iterative process with group discussion and feedback. It was also, however, constrained by data availability. We included covariates representative of potential ecological, environmental, and geomorphological drivers:

##### 1. Vegetation

We used the Normalised Difference Vegetation Index (NDVI) as a proxy for distinguishing vegetation type and the source of SOC. To calculate the NDVI metrics (median and standard deviation), we used Landsat 8 bands from 2014 to 2021 available at 30 m resolution, courtesy of the U.S. Geological Survey, and image processing code from Murray et al. 2022^53,258^ in Google Earth Engine^259^.

##### 2. Elevation and slope

Elevation and slope data was included, as it can be a proxy of soil age and composition, as well as vegetation structure in marshes^260^. Additionally, higher SOC stocks may be associated with shallower slopes, due to lower risk of erosion compared to steeper slopes^261^. We used the Copernicus Digital Elevation Model GLO-30 dataset, which was developed from the TanDEM-X mission between 2011 and 2015. The product is a global dataset of elevation at 30 m resolution and has an absolute vertical accuracy of less than four metres. Slope was then derived using the ee.Terrain.slope() function in Google Earth Engine^259^. The dataset can be found here: https://spacedata.copernicus.eu/collections/copernicus-digital-elevation-model

##### 3. Tidal range

Tidal range can influence the stability and resilience of marshes^262^, as well as the accommodation space^44^. We used the FES2014 Tide Model M2, which has a resolution of ∼7km, available here: https://datastore.cls.fr/catalogues/fes2014-tide-model/

##### 4. Historical relative sea-level rise

Higher SOC stocks further from estuaries can be explained by the signature of past sea level rise^31^. We used the five broad classes originally presented in Clark et al. 1978^39^.

##### 5. Total suspended matter

The Total suspended matter (TSM) was retrieved from the GlobColour project, which processed the TSM data from MERIS imagery collected by the Envisat European Space Agency satellite. Monthly values at 4 km resolution were retrieved from the period 2003-2011, as these were the full years available, which were averaged to generate one TSM layer. The data can be found here: https://hermes.acri.fr/

##### 6. Coastal morphology

Each coastal setting (deltas, estuaries, lagoons, composite deltas and lagoons, bedrock, and carbonate) have an environmental signature that controls the SOC stock via the type and rate of sediment supply to the coastline, nutrient loading/limitation, and organic matter diagenesis^43^. We used the groups of coastlines from the Ecological Coastal Units^263^, which were generated by clustering the 4 million coastal line segments based on 10 variables (two land, five ocean, and three coastline variables). We rasterized the 1 km shoreline segments, which were available from: https://www.arcgis.com/home/item.html?id=54df078334954c5ea6d5e1c34eda2c87

##### 7. Temperature and precipitation

Higher temperatures generally increase the productivity and growth of vegetation^30^, and are associated with higher SOC stocks^29^. Higher rainfall is generally associated with higher SOC by increasing the freshwater runoff and thus potentially higher deposition of allochthonous organic matter^261^. Both minimum and maximum monthly average values of temperature and precipitation were chosen rather than mean annual values^43^, as they portray environmental thresholds that may have a stronger effect on SOC stocks by constraining ecosystem functionality^264^, which regulates both production and decomposition rates. We used data from WorldClim BIO variables^265^ at 927.67 m resolution, available from: https://www.worldclim.org/data/bioclim.html

##### 8. Potential evapotranspiration

Potential evapotranspiration (PET) is the amount of plant evaporation that would occur if there was a sufficient water source in the surrounding soil. This measure has been found to explain different ecophysiological processes in mangroves^266^, and may show a tradeoff in extreme climates. We used the average ENVIREM mean monthly PET of the driest quarter^267^, as this can represent the more extreme cases, when a prolonged dry period can have a negative effect on plant productivity. The dataset can be found here: https://envirem.github.io/#downloads

Most covariate layers (tidal range, TSM, ECUs, temperature, precipitation, PET) were land or ocean products that needed to be extrapolated to each pixel containing tidal marsh. This was done by calculating the average of neighbouring pixels using a circle-shaped boolean kernel in Google Earth Engine^259^, with the functions ee.Image.reduceNeighborhood(), either ee.Reducer.mean() or ee.Reducer.mode() for categorical variables, and ee.Kernel.circle(1, ‘pixels’). To get all covariate map layers to the same 30 m resolution as the DEM and the NDVI layers, we re-sampled and re-projected coarser resolution data to a unified pixel grid (World Geodetic System 1984, EPSG:4326) using bilinear interpolation.

### Model training

We used the random forest model implemented in the ranger package (0.15.1)^268^ and the caret framework (6.0-94)^269^ within R^270^. Due to the limited data in many regions of the world, we used the entire dataset for model training. R^2^ and Root Mean Square Error (RMSE) values were calculated from all pairs of observed and predicted response values when held back from model training using the cross validation method. Variable importance for the random forest model was set to “impurity” within the ranger package, corresponding to the Gini index for classification.

#### k-NNDM cross validation

The choice of a cross-validation strategy is key because it determines the estimation of the model performance as well as the variable importance. Our training data is clustered due to the nature of the available measurements (i.e., large amounts of data in the U.S.A., Australia, and the U.K.), which means that random cross-validation (CV) would only indicate the ability of the model to predict within the clusters^34,271,272^. A spatial cross validation strategy, in which spatial units are held back for validation^272,273^, assesses the ability of the model to predict beyond the clusters, which is in line with the purpose of the model to predict into spaces that lack training data^273–275^. We used the k-fold Nearest Neighbour Distance Matching (k-NNDM) Cross-Validation presented by Linnenbrink et al. 2023^276^ and implemented within the CAST package (0.8.1)^277^, which is a variant of the leave-one-out NNDM cross validation with reduced computation time compared to the method developed by Milà et al. 2022^274^.

This k-NNDM method creates folds such that the geographic distance between sample points of different folds approximates the distance between the training samples and the prediction locations. To create the prediction locations, 5,000 points were randomly selected from the global tidal marsh extent map^4^. We used k = 5 folds (Fig. S6), so that the training data were clustered geographically. When the model separates the training data into training and testing at the cross-validation phase, each one fold serves as testing while the others are used for training. By comparing the geographic distance between folds of our k-NNDM CV to those between random folds, we can see that our method better resembles the distance from prediction locations to training samples (Fig. S7).

#### Model tuning

We implemented a model tuning step, testing 100 possible models by varying the number of variables to consider at each split (1, 2, 3, 4, or 5), the minimum node size (1, 2, 3, 5, or 10), and the number of trees (100, 200, 300, 400, or 500). Between these, the RMSE varied very little, as expected for random forest models^278^. We thus set mtry to 3, the minimum node size to 5, and the number of trees to 300.

### Predictions

To align with the highest resolution covariate variable, the 10 m tidal marsh data was exported at 30 m resolution using Google Earth Engine^259^. The predictions were made for every 30 m pixel identified as tidal marsh in 2020^4^. This recent extent map was derived from earth observation data and estimates 52,880 km^2^ of tidal marshes between 60°N and 60°S. As described in the spatial modelling of soil organic carbon section, the model predicted SOC density at 0 and 30 cm, which were then averaged and multiplied by 30 cm (the layer thickness) and by 100 to get a SOC stock in Mg per hectare. This was also undertaken for the 30-100 cm soil layer, using the SOC density values at 30 and 100 cm. Thus, each 30 m pixel has a predicted SOC stock (Mg ha^-1^) for the 0-30 cm and 30-100 cm soil layers.

### Pixel-wise accuracy estimation

To estimate how different the prediction areas were to areas on which the model was trained, we first calculated a dissimilarity index (DI). In brief, this is calculated by dividing the minimum distance to the nearest training data point in a multidimensional predictor space that has been scaled and weighted by variable importance, by the average of the distances in the training data^45^. The DI is calculated based on data points that do not occur in the same cross-validation fold, thus keeping in mind the cross-validation nature of the model.

Then, we used the relationship between the DI and the prediction performance (i.e. the final model RMSE) to produce a spatially continuous estimation of the expected error associated with each SOC prediction (DItoErrormetric() in developer’s CAST commit version from August 2023)^279^, This uses shape constrained additive models^280^ to model the relationship between the DI and the RMSE. This model can then be applied to the DI of every tidal marsh pixel to produce a spatially continuous map of the estimated accuracy of the predictions. We calculated the expected model error similarly as we did for the predictions using the spatial modelling approach, and to get an expected error in the same units as the predictions.

The analysis workflow (model training, predictions, error, and AOA) was completed using Snakemake^281^. We calculated average prediction and expected error values for eleven out of the twelve biogeographical realms of the Marine Ecoregions of the World^33^ (Fig. 3, Table S3). There was no tidal marsh extent predicted in the twelve realm, Southern Ocean. We also calculated averages per country using the union of the ESRI Country shapefile and the Exclusive Economic Zones from the Flanders Marine Institute (2020)^282^.

### Area of applicability

Although we can apply the Random Forest model to all tidal marshes globally because of the availability and preparation of covariate data for all marshes, these predictions can often be extrapolated and rendered meaningless when predictor values are too different compared to the training data^275^. We thus implemented the area of applicability (AOA), introduced by Meyer and Pebesma 2021^45^, to mask out areas where the model wasn’t able to learn about the relationship between the predictors and the response (here, soil organic carbon density).

The threshold for the AOA is determined by the outlier-removed maximum DI of the training data, i.e. data larger than the 75th percentile plus 1.5 times the interquartile range of the DI values of the cross-validated training data. The calculation of the pixel-level DI and the AOA were generated using the aoa() function, both available in the CAST package (0.8.1)^277^. Then, the predictor space which is greater than the AOA threshold is considered outside the area of applicability, and thus is masked from our predictions. For each predicted soil layer (i.e. 0-30 cm and 30-100 cm), the AOA was calculated at the upper and lower depth and then averaged. Locations considered inside the AOA (Fig. S1b & Fig. S2b) are those where the averaged AOA is equal to 1 (i.e. AOA values of 0 or 0.5 are considered outside the AOA).

## Data availability

The training data, tidal marsh extent and environmental covariate data used in this study are publicly available (linked in the Methods). The soil organic carbon predictions, estimated model errors, and area of applicability layer for both the 0-30 cm and 30-100 cm layers are available on Zenodo (https://doi.org/10.5281/zenodo.10940066).

## Code availability

All code used in this study is available on Github: https://github.com/Tania-Maxwell/global-marshC-map/

## Acknowledgements

We thank Daniele Baisero, Thomas Ball, and Alison Eyres for methodological help. This project benefited from funding from the Bezos Earth Fund and other donors supporting the Nature Conservancy. LH Pérez-Bernal provided assistance in geochemical analysis of sediment cores from Mexico. This work was performed using resources provided by the Cambridge Service for Data Driven Discovery (CSD3) operated by the University of Cambridge Research Computing Service (www.csd3.cam.ac.uk), provided by Dell EMC and Intel using Tier-2 funding from the Engineering and Physical Sciences Research Council (capital grant EP/T022159/1), and DiRAC funding from the Science and Technology Facilities Council (www.dirac.ac.uk). Any use of trade, firm, or product names is for descriptive purposes only and does not imply endorsement by the U.S. Government.

## Author information

### Contributions

T.A.W., M.D.S., E.L., and T.L.M. and designed the research. T.L.M. and T.A.W. performed the analyses. M.F.A., J.A., M.Cop., M.Cos, G.C., D.F., J.H., C.La., C.Lo, M.M.M., J.R., K.R.,

A.C.R.-F., O.S., C.S., M.V.d.B., and L.W.-M. provided expert knowledge on tidal marsh soil carbon dynamics. C.La. M.L., N.M., A.N., A.R., and L.W. helped with the methodology. E.L. provided funding. M.D.S. L.S.S. and T.A.W. provided supervision. The manuscript was drafted by T.L.M. and T.A.W. with contributions from all co-authors.

## Ethics declarations

The authors declare no competing interests.

