## Supplementary Information for "Soil carbon in the world’s tidal marshes"

Accompanying article: Maxwell et al. (2024) Soil carbon in the world's tidal marshes.

### Figures

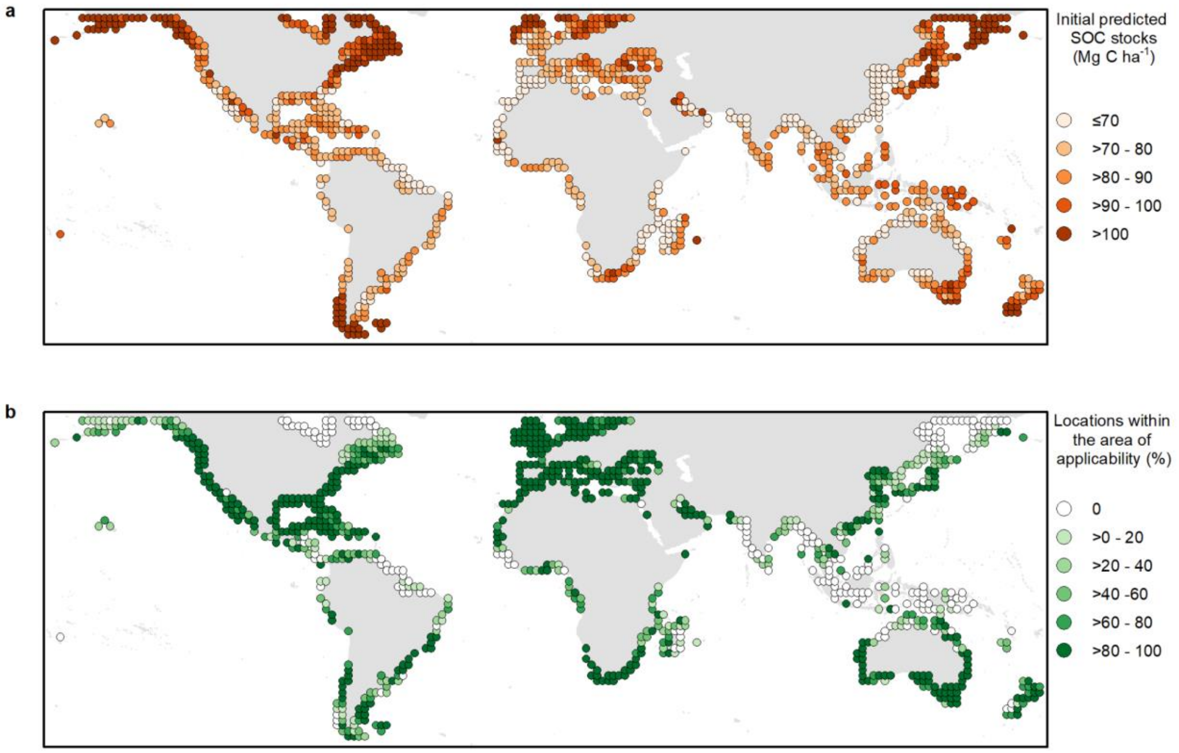


**Fig. S1 | Global distribution of tidal marsh soil organic carbon (SOC) for the 0-30 cm soil layer (aggregated per 2° cell).** a) Initial predicted SOC per unit area (Mg C ha$^{-1}$). b) The proportion of pixels located within the area of applicability (AOA), i.e. where we enabled the model to learn about the relationship between SOC and the environmental drivers for this 0-30 cm soil layer. The final predicted SOC per unit area (Mg C ha$^{-1}$), after removing pixels outside the AOA, are presented in Fig. 2a.

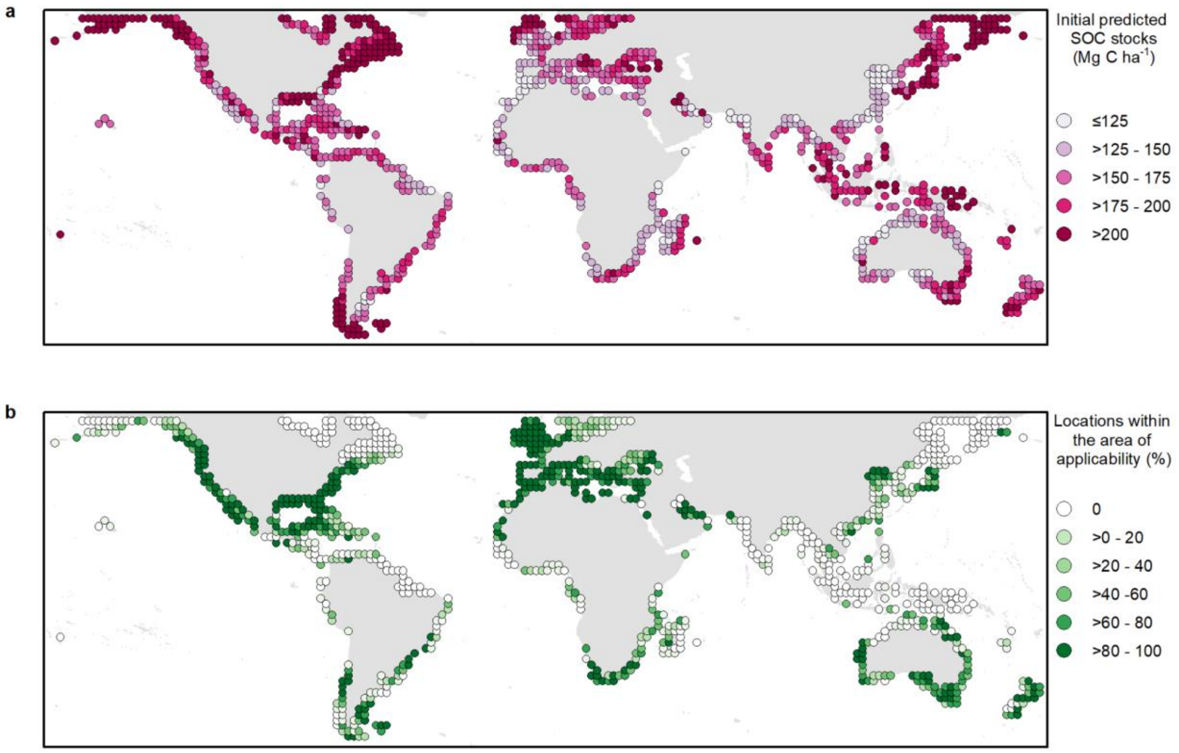


**Fig. S2 | Global distribution of tidal marsh soil organic carbon (SOC) for the 30-100 cm soil layer (summarised per 2° cell).** a) Initial predicted SOC per unit area (Mg C $ha^{-1}$). b) The proportion of pixels located within the area of applicability (AOA), i.e. where we enabled the model to learn about the relationship between SOC and the environmental drivers for this 30-100 cm soil layer. The final predicted SOC per unit area (Mg C $ha^{-1}$), after removing pixels outside the AOA, are presented in Fig. 2b.

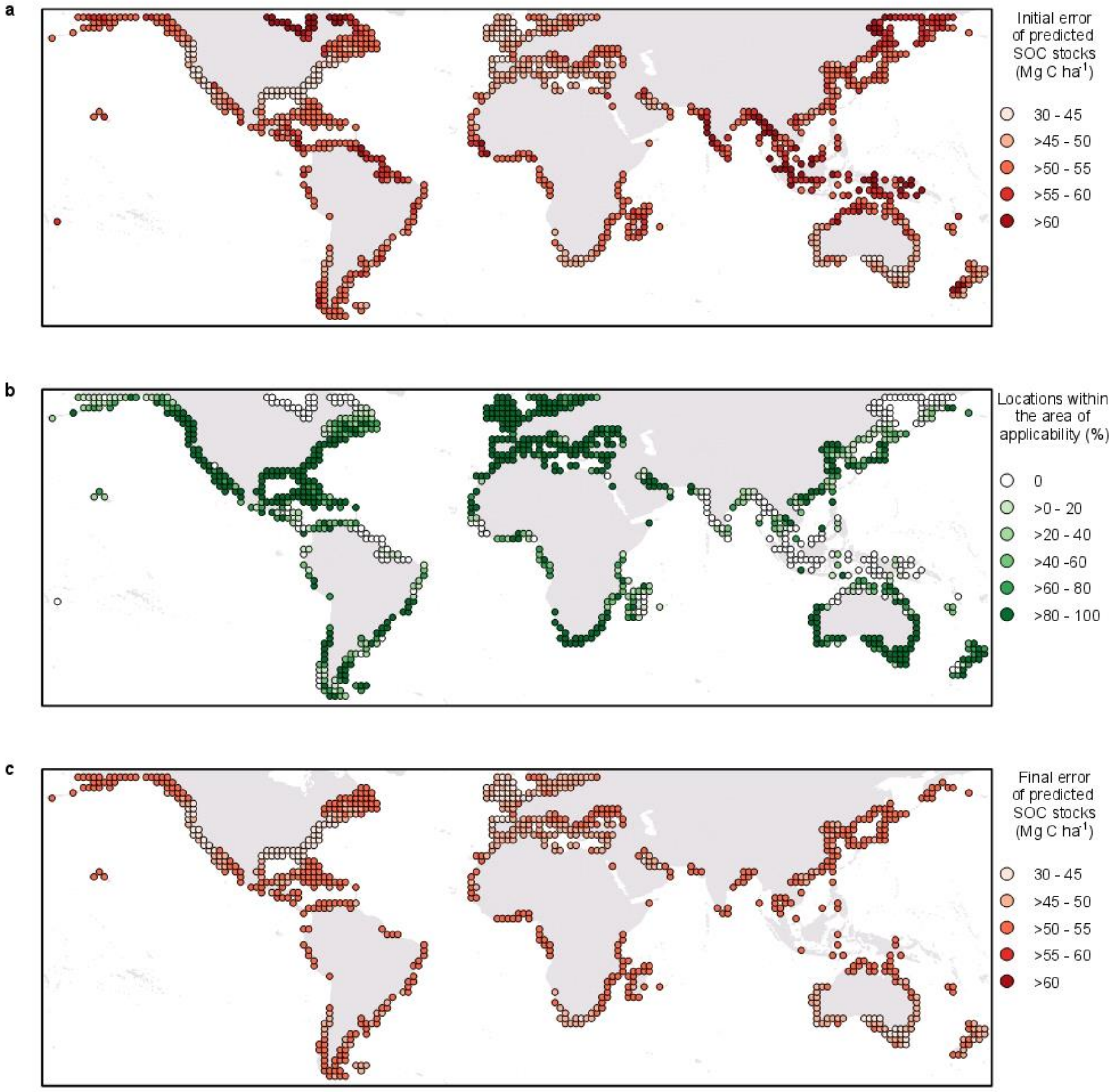


**Fig. S3 |** Global distribution of expected error of the tidal marsh soil organic carbon (SOC) predictions for the 0-30 cm soil layer (aggregated to 2°). a) Initial expected model error of predicted SOC per unit area (Mg C $ha^{-1}$), without taking into account whether predictions were meaningful. b) The proportion of pixels located within the area of applicability (AOA), i.e. where we enabled the model to learn about the relationship between SOC and the environmental drivers for this 0-30 cm soil layer. c) Final expected model error of predicted SOC per unit area (Mg C $ha^{-1}$), after masking out pixels outside the AOA.

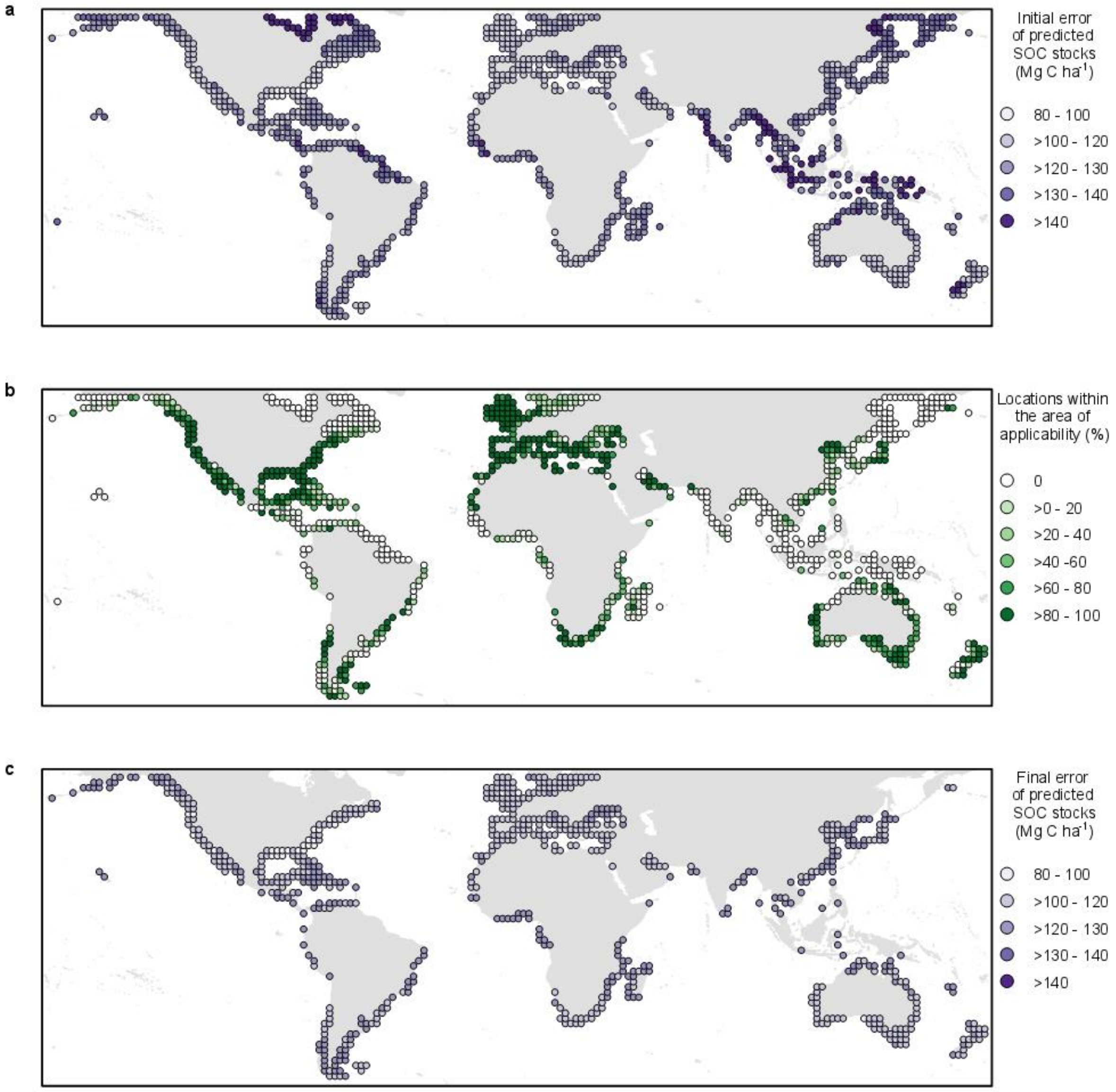


**Fig. S4 |** Global distribution of expected error of the tidal marsh soil organic carbon (SOC) predictions for the 30-100 cm soil layer (summarised to 2°). a) Initial expected model error of predicted SOC per unit area (Mg C ha$^{-1}$), without taking into account whether predictions were meaningful. b) The proportion of pixels located within the area of applicability (AOA), i.e. where we enabled the model to learn about the relationship between SOC and the environmental drivers for this 0-30 cm soil layer. c) Final expected model error of predicted SOC per unit area (Mg C ha$^{-1}$), after masking out pixels outside the AOA.

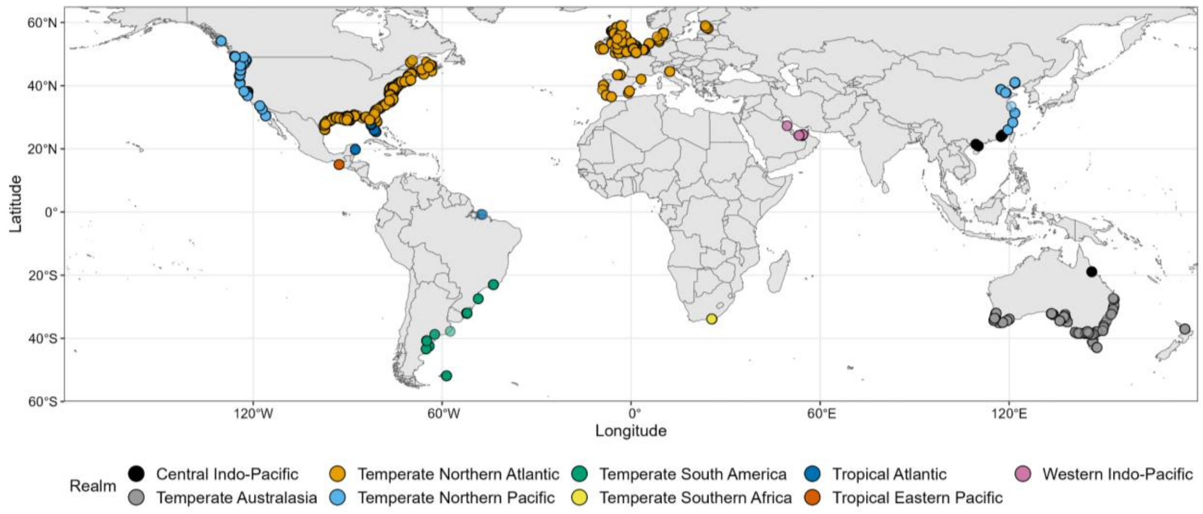


**Fig. S5 |** Locations of training data points for the biogeographical realms of the Marine Ecoregions of the World[33]. The arctic and eastern Indo-Pacific realms are not represented due to lack of data in these regions.

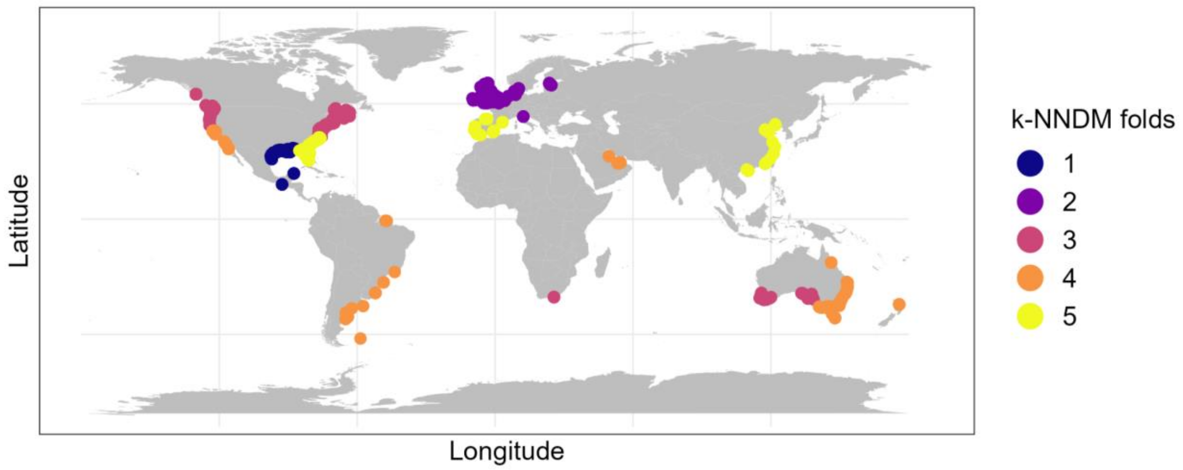


**Fig. S6 |** Training data divided into 5 folds for the k-fold Nearest Neighbour Distance Matching (k-NNDM) Cross-Validation, following methods described by Linnenbrink et al. 2023[73].

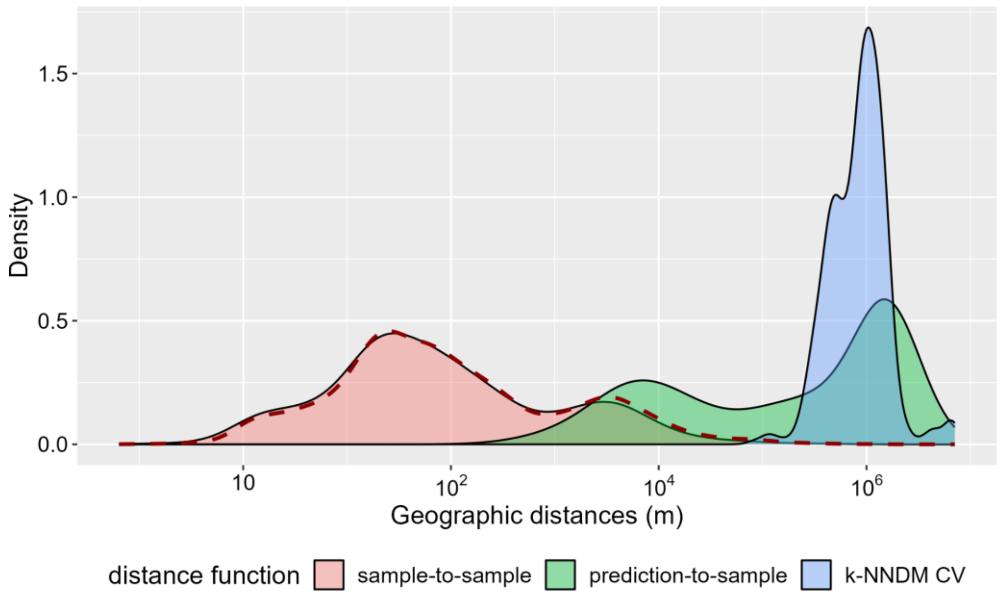


**Fig. S7 |** Comparison of the geographic distance between folds of the random cross validation (dashed red line), which reproduces the distances between the samples (pink), and the k-fold nearest neighbour distance matching (k-NNDM) cross validation (blue), which better resembles the distance from prediction locations to training samples (green).

#### Tables

**Table S1.** Hypothesized landscape-level drivers of soil organic carbon (SOC) in tidal marshes globally. These variables were based on the literature, along with previous studies investigating the variables identified for their associations with SOC in vegetated coastal ecosystems[15,283,284].

| **Landscape-level drivers** | **Relation to SOC in tidal marshes** | **Spatial data description** | **Spatial data source** |
|---|---|---|---|
| ECOLOGICAL | | | |
| Vegetation class | C stocks vary across species [41,42]. Species with rhizomes (graminoids) have higher SOC density. | Use NDVI as a proxy for source of SOC. Calculated from Landsat 8 bands from 2014 to 2021.<br><br>→ resolution 30 m | ee.ImageCollection("LANDSAT/LC08/C02/T1_L2")<br><br>Pre-processed using code from Murray et al. (2022)[53,258] |
| GEOMORPHOLOGICAL | | | |
| Elevation (m) | Higher SOC stocks are generally found in lower elevation due (1) higher sedimentation rates allowing more trapping of organic C from surface organisms[36] and (2) more frequent inundation providing opportunity to settle more allochthonous C particles[37]. However, sea level rise history may lead to higher C stocks in the intertidal marshes[31,285]. | Copernicus DEM<br>→ resolution 30 m | ee.ImageCollection("COPERNICUS/DEM/GLO30")<br>https://spacedata.copernicus.eu/documents/20123/121239/GEO1988-CopernicusDEM-SPE-002_ProductHandbook_I4.0.pdf |

| | | | |
|---|---|---|---|
| Slope (%) | Higher SOC stocks are associated with shallower slopes, due to lower risk of erosion compared to steeper slopes[261]. | Calculated from elevation data | |
| Tidal range (m) | Can influence the stability and resilience of marshes[262], as well as accommodation space[44]. | FES2014 Tide Model M2<br>→ resolution 1/16 degree (~7 km) | https://datastore.cls.fr/catalogues/fes2014-tide-model/ |
| Sea-level rise | Higher SOC stocks further from estuaries can be explained by the signature of past sea level rise[31]. | Holocene relative sea-level rise zones<br>→ 5 broad-scale zones | Clark et al. (1987)[39] |
| Coastal morphology | Each coastal setting (deltas, estuaries, lagoons, composite deltas and lagoons, bedrock, and carbonate) has an apparent environmental signature that controls the SOC stock *via* the type and rate of sediment supply to the coastline, nutrient loading/limitation, and organic matter diagenesis[43]. | Ecological coastal units<br>→ 16 classes | https://www.esri.com/arcgis-blog/products/arcgis-living-atlas/mapping/ecus-available/ |
| OTHER ENVIRONMENTAL | | | |
| Minimum temperature of coldest month and maximum temperature of warmest month (°C) | Higher temperatures generally increase the productivity and growth of vegetation[30], and are associated with higher SOC stocks[29].<br><br>Minimum temperature and minimum precipitation were chosen rather than mean annual values[43], as they portray environmental thresholds that may have a stronger effect on SOC stocks by constraining ecosystem functionality[264], which regulates both production and decomposition rates. | WorldClim BIO6<br>→ resolution 927.67 meters | https://www.worldclim.org/data/bioclim.html<br><br>ee.Image("WORLDCLIM/V1/BIO") |
| Precipitation of driest month (mm) and precipitation of wettest month (mm) | Higher rainfall is generally associated with higher SOC by increasing the freshwater runoff and thus potentially higher deposition of allochthonous organic matter[261]. | WorldClim BIO14<br>→ resolution 927.67 meters | https://www.worldclim.org/data/bioclim.html<br><br>ee.Image("WORLDCLIM/V1/BIO") |
| Potential evapotranspiration (PET) of the driest quarter and PET of the warmest quarter (mm / month) | Potential evapotranspiration has been found to explain ecophysiological processes in mangroves, a neighbouring coastal ecosystem. The potential evapotranspiration can influence the concentration of salts and nutrients in the soils and groundwater in wetlands[286]. | ENVIREM<br>→ resolution 927.67 meters | https://envirem.github.io/#download |
| Total suspended matter | More sediment is associated with lower C density but deeper deposits, along with higher accretion and C sequestration[287]. | Derived from remotely sensed MERIS, reprocessed from GlobColour project<br>→ Resolution 4 km<br>→ Monthly values (2003-2011) averaged per quarter[35] | https://hermes.acri.fr/ |

**Table S2 | Country-level summary statistics for the tidal marsh global soil organic carbon (SOC) map.** For each soil layer (0-30 cm and 30-100 cm), the initial predicted SOC stock, the proportion of the realm within the area of applicability (AOA), and the final predicted SOC stock, after masking out areas outside the AOA. The expected model error is shown in parentheses for each prediction. Only countries with a tidal marsh extent greater than 10 km$^2$ are represented here.

| | 0-30 cm | | | 30-100 cm | | | | |
|---|---|---|---|---|---|---|---|---|
| **Country** | **Initial predicted SOC stock (Mg ha$^{-1}$)** | **Locations within the area of applicability (%)** | **Final predicted SOC stock (Mg ha$^{-1}$)** | **Initial predicted SOC stock (Mg ha$^{-1}$)** | **Locations within the area of applicability (%)** | **Final predicted SOC stock (Mg ha$^{-1}$)** | **Area (km$^2$)** | **Total C to 1 m (Tg)** |
| Albania | 80.38 (50.14) | 94.8 | 79.79 (49.87) | 169.69 (119.07) | 82.2 | 162.90 (117.60) | 79.01 | 1.92 (1.32) |
| Algeria | 69.70 (51.29) | 78.5 | 68.98 (50.1) | 148.01 (121.23) | 67.4 | 142.25 (117.78) | 26.92 | 0.57 (0.45) |
| Angola | 71.06 (53.22) | 68.5 | 69.49 (52.75) | 145.73 (125.95) | 0 | 141.75 (123.97) | 17.04 | 0.36 (0.30) |
| Argentina | 78.84 (53.28) | 43.4 | 77.46 (51.73) | 166.46 (126.78) | 9.2 | 166.75 (121.04) | 2429.48 | 59.33 (41.97) |
| Australia | 86.12 (47.41) | 82.8 | 88.08 (45.69) | 170.04 (116.91) | 64.7 | 172.86 (110.87) | 2080.51 | 54.29 (32.57) |
| Bahamas | 73.89 (51.56) | 80.7 | 74.24 (51.16) | 156.85 (123.63) | 39.3 | 154.76 (120.45) | 116.29 | 2.66 (2.00) |
| Bangladesh | 54.59 (58.04) | 9.3 | 56.80 (52.81) | 118.82 (137.07) | 0.7 | 127.67 (122.04) | 202.47 | 3.74 (3.54) |
| Belize | 77.53 (49.18) | 81.1 | 76.67 (47.71) | 163.39 (118.6) | 73.7 | 159.31 (115.09) | 64.17 | 1.51 (1.04) |
| Benin | 78.45 (52.83) | 70.9 | 78.53 (52.33) | 165.67 (124.67) | 5.7 | 168.67 (122.27) | 53.94 | 1.33 (0.94) |
| Brazil | 73.61 (52.52) | 58.3 | 76.11 (49.4) | 164.26 (126.19) | 34.7 | 171.06 (117.32) | 1066.11 | 26.35 (17.77) |
| Bulgaria | 96.09 (52.15) | 93.2 | 95.48 (52.01) | 200.18 (122.97) | 58.5 | 193.87 (121.78) | 19.55 | 0.57 (0.34) |
| Cambodia | 79.76 (52.62) | 76.7 | 80.04 (52.14) | 165.66 (124.28) | 34.4 | 163.94 (121.92) | 15.8 | 0.39 (0.28) |
| Canada | 94.11 (60.56) | 8.5 | 104.97 (49.03) | 186.27 (143.71) | 3.2 | 227.79 (115.79) | 8534.32 | 283.99 (140.66) |
| Chile | 116.86 (54.01) | 39.6 | 103.14 (49.97) | 234.62 (127.9) | 33.4 | 201.6 (118.49) | 458.62 | 13.98 (7.73) |
| China | 57.43 (50.88) | 80.6 | 57.28 (49.97) | 124.07 (121.81) | 34.7 | 108.41 (117.65) | 1167.51 | 19.34 (19.57) |
| Colombia | 77.11 (53.58) | 50 | 77.55 (52.03) | 169.39 (126.63) | 18.3 | 171.12 (121.27) | 45.93 | 1.14 (0.80) |
| Costa Rica | 71.7 (53.55) | 38.4 | 71.98 (53.06) | 148.61 (125.81) | 2.6 | 149.71 (122.89) | 15.00 | 0.33 (0.26) |
| Croatia | 93.53 (52.30) | 61.8 | 91.66 (51.24) | 202.58 (124.31) | 29.6 | 193.42 (119.34) | 51.76 | 1.48 (0.88) |
| Cuba | 85.10 (50.53) | 97.4 | 85.22 (50.44) | 178.55 (119.92) | 90.9 | 179.13 (119.44) | 783.79 | 20.72 (13.32) |
| Denmark | 87.45 (48.54) | 91.3 | 86.56 (47.93) | 171.42 (118.18) | 48.6 | 162.38 (113.05) | 263.73 | 6.57 (4.25) |
| Dominican Republic | 88.12 (51.91) | 82.2 | 86.42 (51.4) | 195.03 (123.26) | 41.8 | 174.84 (120.22) | 17.37 | 0.45 (0.30) |
| Egypt | 71.26 (49.40) | 94.8 | 70.60 (49.12) | 146.63 (117.09) | 92.3 | 144.07 (116.26) | 176.83 | 3.80 (2.92) |
| Estonia | 114.09 (45.82) | 93.1 | 113.74 (45.09) | 219.54 (116.9) | 23.4 | 211.06 (113.97) | 269.45 | 8.75 (4.29) |
| France | 78.41 (49.83) | 89.8 | 77.83 (49.23) | 162.98 (118.64) | 81.6 | 158.25 (116.74) | 678.47 | 16.02 (11.26) |
| Gabon | 77.45 (53.28) | 65.6 | 76.30 (52.27) | 162.03 (125.04) | 53.6 | 159.02 (122.01) | 34.69 | 0.82 (0.60) |
| Gambia | 63.29 (53.95) | 24.3 | 61.6 (53.00) | 132 (127.25) | 0.3 | 145.79 (122.13) | 40.5 | 0.84 (0.71) |

| | | | | | | | | |
|---|---|---|---|---|---|---|---|---|
| Georgia | 89.32 (51.39) | 86.1 | 88.01 (50.93) | 234.36 (121.84) | 63.2 | 232.56 (119.57) | 65.04 | 2.08 (1.11) |
| Germany | 78.36 (46.39) | 97.6 | 77.71 (46.19) | 157.88 (113.02) | 71.9 | 143.10 (109.33) | 346.03 | 7.64 (5.38) |
| Ghana | 71.94 (52.63) | 90.1 | 70.95 (52.46) | 145.83 (124.38) | 5.8 | 146.91 (122.5) | 16.96 | 0.37 (0.30) |
| Greece | 81.27 (49.85) | 93.6 | 80.75 (49.57) | 170.87 (118.55) | 83.2 | 166.81 (117.18) | 213.35 | 5.28 (3.56) |
| Guatemala | 90.00 (53.03) | 62.1 | 92.11 (51.98) | 192.07 (125.12) | 33.8 | 199.25 (121.86) | 11.56 | 0.34 (0.20) |
| Guinea-Bissau | 63.46 (57.03) | 10.0 | 59.19 (52.23) | 136.49 (134.55) | 2.2 | 125.01 (121.68) | 68.82 | 1.27 (1.2) |
| Guyana | 74.00 (59.96) | 0 | NA | 170.37 (141.28) | 0 | NA | 33.28 | NA |
| Haiti | 73.48 (53.70) | 42.8 | 72.45 (52.39) | 157.60 (127.00) | 4.1 | 184.12 (119.60) | 38.64 | 0.99 (0.66) |
| Honduras | 81.24 (54.44) | 18.7 | 74.83 (52.92) | 175.59 (128.01) | 0.3 | 150.88 (121.36) | 393.12 | 8.87 (6.85) |
| India | 64.84 (55.75) | 31.1 | 69.72 (52.13) | 139.07 (132.23) | 8.1 | 153.00 (120.2) | 52.3 | 1.16 (0.9) |
| Indonesia | 75.82 (59.06) | 0.4 | 86.01 (52.32) | 178.56 (139.57) | 0.1 | 180.18 (122.09) | 170.73 | 4.54 (2.98) |
| Iran | 92.16 (54.00) | 16.4 | 64.41 (46.96) | 209.04 (127.41) | 15.5 | 128.88 (114.06) | 11.10 | 0.21 (0.18) |
| Ireland | 88.31 (45.36) | 96.4 | 87.56 (45.06) | 165.85 (108.88) | 90.4 | 162.14 (106.94) | 130.69 | 3.26 (1.99) |
| Italy | 78.54 (48.43) | 97.0 | 77.40 (48.24) | 164.25 (115.97) | 89.3 | 160.04 (114.87) | 228.72 | 5.43 (3.73) |
| Jamaica | 83.31 (51.69) | 97.4 | 83.21 (51.64) | 176.45 (122.59) | 51.5 | 176.65 (121.61) | 20.68 | 0.54 (0.36) |
| Japan | 86.14 (54.01) | 43.0 | 84.16 (51.62) | 180.90 (128.61) | 4.2 | 212.61 (119.15) | 267.64 | 7.94 (4.57) |
| Latvia | 95.12 (49.23) | 89.5 | 94.87 (48.44) | 184.26 (120.88) | 25.0 | 177.95 (117.04) | 163.61 | 4.46 (2.71) |
| Lithuania | 87.18 (49.67) | 85.7 | 86.73 (48.77) | 170.07 (121.39) | 8.4 | 168.26 (117.76) | 37.97 | 0.97 (0.63) |
| Madagascar | 71.52 (53.96) | 37.6 | 65.06 (51.27) | 152.57 (127.15) | 26.2 | 131.66 (119.89) | 153.32 | 3.02 (2.62) |
| Mauritania | 70.38 (50.08) | 98.8 | 70.45 (50.03) | 139.34 (119.85) | 68.0 | 144.5 (118.18) | 28.26 | 0.61 (0.48) |
| Mexico | 79.68 (49.59) | 90.7 | 79.25 (49.06) | 171.64 (118.79) | 79.6 | 169.13 (116.82) | 1194.69 | 29.67 (19.82) |
| Montenegro | 90.85 (54.28) | 31.4 | 92.88 (50.81) | 194.46 (129.49) | 18.1 | 192.47 (118.95) | 26.66 | 0.76 (0.45) |
| Morocco | 59.22 (48.77) | 87.5 | 58.37 (48.05) | 114.95 (116.53) | 75 | 108.94 (113.35) | 43.02 | 0.72 (0.69) |
| Mozambique | 63.81 (52.96) | 55.8 | 63.72 (51.70) | 135.42 (125.15) | 29.5 | 131.52 (121.00) | 1343.19 | 26.22 (23.20) |
| Myanmar | 59.41 (68.44) | 0 | NA | 129.19 (162.63) | 0 | NA | 122.99 | NA |
| Netherlands | 82.41 (47.24) | 86.5 | 75 (45.8) | 181.83 (114.78) | 79.1 | 160.04 (110.98) | 183.33 | 4.31 (2.87) |
| New Zealand | 92.06 (52.51) | 70.0 | 90.14 (49.46) | 183.97 (125.09) | 53.9 | 175.11 (116.50) | 244.82 | 6.49 (4.06) |
| Nicaragua | 76.51 (54.95) | 12.9 | 79.17 (52.99) | 162.46 (128.80) | 8.5 | 162.68 (123.64) | 559.72 | 13.54 (9.89) |
| Nigeria | 78.18 (54.01) | 31.7 | 79.77 (52.82) | 169.71 (127.22) | 3.8 | 157.82 (123.56) | 21.42 | 0.51 (0.38) |
| North Korea | 63.56 (53.57) | 41.4 | 58.94 (52.26) | 132.06 (125.35) | 35.6 | 122.07 (121.92) | 80.01 | 1.45 (1.39) |
| Papua New Guinea | 81.76 (58.32) | 3.0 | 88.16 (52.82) | 185.78 (138.09) | 0.2 | 186.95 (122.52) | 53.25 | 1.46 (0.93) |
| Peru | 72.84 (52.37) | 68.9 | 71.28 (51.43) | 145.87 (125.54) | 13.0 | 129.47 (120.02) | 10.08 | 0.20 (0.17) |
| Poland | 95.84 (49.29) | 95.3 | 95.28 (49.03) | 183.46 (120.51) | 20.0 | 179.49 (118.04) | 161.17 | 4.43 (2.69) |
| Portugal | 58.56 | 97.6 | 58.33 | 109.84 | 96.3 | 108.87 | 157.7 | 2.64 |

| | | | | | | | | |
|---|---|---|---|---|---|---|---|---|
| | (44.17) | | (43.89) | (108.51) | | (107.71) | | (2.39) |
| Romania | 91.26 (51.93) | 97.3 | 91.21 (51.88) | 187.77 (123.24) | 28.7 | 187.15 (122.28) | 667.23 | 18.57 (11.62) |
| Russia | 106.10 (56.11) | 26.2 | 91.10 (51.04) | 208.2 (132.76) | 20.8 | 180.12 (119.29) | 5140.54 | 139.42 (87.56) |
| Senegal | 63.90 (51.45) | 95.5 | 63.93 (51.33) | 127.10 (122.15) | 37.0 | 126.67 (120.94) | 52.43 | 1.00 (0.9) |
| Sierra Leone | 67.96 (57.92) | 0 | NA | 141.94 (136.25) | 0 | NA | 21.08 | NA |
| South Africa | 82.69 (47.23) | 96.6 | 82.32 (46.97) | 165.13 (114.6) | 81.0 | 158.85 (112.5) | 89.15 | 2.15 (1.42) |
| South Korea | 61.36 (54.42) | 18.3 | 63.12 (52.85) | 125.03 (127.32) | 6.6 | 114.86 (123.52) | 180.83 | 3.22 (3.19) |
| Spain | 62.07 (46.21) | 98.3 | 61.99 (46.05) | 117.78 (111.96) | 95.8 | 116.61 (111.3) | 341.47 | 6.10 (5.37) |
| Suriname | 69.21 (57.06) | 0 | NA | 157.88 (134.34) | 0 | NA | 42.44 | NA |
| Sweden | 94.47 (49.54) | 87.9 | 93.3 (48.69) | 178.61 (120.88) | 31.5 | 164.59 (115.81) | 117.08 | 3.02 (1.93) |
| Thailand | 82.76 (54.15) | 40 | 86.4 (51.92) | 180.60 (127.74) | 21 | 185.52 (121.33) | 18.25 | 0.50 (0.32) |
| Tunisia | 66.59 (46.31) | 98.5 | 65.53 (46.15) | 134.90 (109.91) | 97.8 | 131.81 (109.43) | 53.80 | 1.06 (0.84) |
| Turkey | 88.35 (49.08) | 94 | 88.35 (48.75) | 189.20 (117.65) | 82.9 | 187.61 (116) | 252.44 | 6.97 (4.16) |
| Ukraine | 87.80 (52.14) | 87.4 | 87.64 (51.76) | 175.85 (123.52) | 38.7 | 172.70 (121.95) | 923.82 | 24.05 (16.05) |
| United Kingdom | 88.53 (42.84) | 98 | 87.97 (42.6) | 171.60 (105.58) | 93.2 | 167.37 (104.12) | 535.86 | 13.68 (7.86) |
| United States | 86.71 (42.45) | 84.4 | 82.76 (39.99) | 201.63 (104.27) | 82 | 198.86 (98.54) | 18509.72 | 521.27 (256.42) |
| Uruguay | 83.39 (51.17) | 85.1 | 83.02 (50.5) | 185.71 (122.46) | 50.6 | 183.21 (119.55) | 286.62 | 7.63 (4.87) |
| Venezuela | 79.24 (54.59) | 30.1 | 80.62 (51.32) | 174.25 (129.38) | 13.1 | 178.49 (119.04) | 165.20 | 4.28 (2.81) |
| Vietnam | 77.34 (54.97) | 36.8 | 81.24 (52.51) | 165.93 (129.91) | 13.4 | 166.02 (122.72) | 12.08 | 0.30 (0.21) |

**Table S3 | Realm level summary statistics for the tidal marsh global soil organic carbon (SOC) map.** For each soil layer (0-30 cm and 30-100 cm), we present the initial predicted SOC stock, the proportion of the realm within the area of applicability (AOA), i.e. where we enabled the model to learn about the relationship between SOC stocks and the environmental drivers, and the final predicted SOC stock, after masking out areas outside the AOA. The estimated model error is shown in parentheses for each prediction. Realms correspond to the biogeographical realms of the Marine Ecoregions of the World[33].

| | **0-30 cm** | | | **30-100 cm** | | |
|---|---|---|---|---|---|---|
| **Realm** | **Initial predicted SOC stock (Mg ha$^{-1}$)** | **Pixels within the area of applicability (%)** | **Final predicted SOC stock (Mg ha$^{-1}$)** | **Initial predicted SOC stock (Mg ha$^{-1}$)** | **Pixels within the area of applicability (%)** | **Final predicted SOC stock (Mg ha$^{-1}$)** |
| Arctic | 96.19 (60.55) | 1.5 | 121.80 (52.49) | 187.67 (143.15) | 0.3 | 252.54 (122.05) |
| Central Indo-Pacific | 70.55 (56.3) | 22.6 | 71.35 (51.09) | 155.06 (133.03) | 13.4 | 149.19 (118.86) |
| Eastern Indo-Pacific | 82.06 (54.45) | 31.3 | 83.88 (52.44) | 167.15 (128.98) | 5.0 | 188.53 (120.9) |
| Temperate Australasia | 89.56 (47.10) | 87.9 | 89 (45.87) | 176.2 (116.30) | 68.9 | 174.04 (111.19) |
| Temperate Northern Atlantic | 84.95 (43.56) | 95.2 | 84.24 (42.98) | 192.62 (106.58) | 81.8 | 190.87 (102.60) |
| Temperate Northern Pacific | 98.34 (54.80) | 29.2 | 77.11 (48.19) | 196.24 (130.45) | 17.7 | 161.19 (112.22) |
| Temperate South America | 84.41 (52.75) | 52.0 | 81.16 (50.75) | 178.87 (125.75) | 22.4 | 180.69 (118.75) |
| Temperate Southern Africa | 83.31 (47.13) | 96.4 | 82.96 (46.84) | 165.96 (114.48) | 78.8 | 159.59 (112.08) |
| Tropical Atlantic | 77.52 (51.65) | 65.4 | 78.77 (49.46) | 166.04 (123.30) | 53.4 | 168.92 (117.76) |
| Tropical Eastern Pacific | 82.72 (50.44) | 67.7 | 86.42 (48.5) | 178.76 (120.81) | 45.6 | 195.89 (113.88) |
| Western Indo-Pacific | 63.56 (54.63) | 44.9 | 63.95 (51.58) | 135.55 (129.16) | 24.1 | 131.97 (120.60) |
| *Global* | 88.13 (50.63) | 58.0 | 83.09 (44.77) | 186.60 (121.69) | 46.2 | 185.27 (105.71) |

#### All references


1. Lovelock, C. E. & Duarte, C. M. Dimensions of Blue Carbon and emerging perspectives. *Biol. Lett.* **15**, 20180781 (2019).
2. Macreadie, P. I. *et al.* The future of Blue Carbon science. *Nat. Commun.* **10**, (2019).
3. Ouyang, X. & Lee, S. Y. Updated estimates of carbon accumulation rates in coastal marsh sediments. *Biogeosciences* **11**, 5057–5071 (2014).
4. Worthington, T. A. *et al.* The distribution of global tidal marshes from earth observation data. 2023.05.26.542433 Preprint at https://doi.org/10.1101/2023.05.26.542433 (2023).
5. Davidson, N. C. How much wetland has the world lost? Long-term and recent trends in global wetland area. *Mar. Freshw. Res.* **65**, 934 (2014).
6. Mcleod, E. *et al.* A blueprint for blue carbon: toward an improved understanding of the role of vegetated coastal habitats in sequestering CO2. *Front. Ecol. Environ.* **9**, 552–560 (2011).
7. Macreadie, P. I., Allen, K., Kelaher, B. P., Ralph, P. J. & Skilbeck, C. G. Paleoreconstruction of estuarine sediments reveal human-induced weakening of coastal carbon sinks. *Glob. Change Biol.* **18**, 891–901 (2012).
8. Campbell, A. D., Fatoyinbo, L., Goldberg, L. & Lagomasino, D. Global hotspots of salt marsh change and carbon emissions. *Nature* **612**, 701–706 (2022).
9. Hopkinson, C. S., Cai, W.-J. & Hu, X. Carbon sequestration in wetland dominated coastal systems—a global sink of rapidly diminishing magnitude. *Curr. Opin. Environ. Sustain.* **4**, 186–194 (2012).
10. Lotze, H. K. *et al.* Depletion, Degradation, and Recovery Potential of Estuaries and Coastal Seas. *Science* **312**, 1806–1809 (2006).
11. McMahon, L. *et al.* Maximizing blue carbon stocks through saltmarsh restoration. *Front. Mar. Sci.* **10**, (2023).
12. Granek, E. F. *et al.* Ecosystem Services as a Common Language for Coastal Ecosystem-Based Management. *Conserv. Biol.* **24**, 207–216 (2010).
13. Holmquist, J. R. *et al.* Accuracy and Precision of Tidal Wetland Soil Carbon Mapping in the Conterminous United States. *Sci. Rep.* **8**, 9478 (2018).
14. Smeaton, C. *et al.* Using citizen science to estimate surficial soil Blue Carbon stocks in Great British saltmarshes. *Front. Mar. Sci.* **9**, 959459 (2022).
15. Young, M. A. *et al.* National scale predictions of contemporary and future blue carbon storage. *Sci. Total Environ.* **800**, (2021).
16. Macreadie, P. I. *et al.* Blue carbon as a natural climate solution. *Nat. Rev. Earth Environ.* **2**, 826–839 (2021).
17. Hengl, T. *et al.* SoilGrids250m: Global gridded soil information based on machine learning. *PLOS ONE* **12**, e0169748 (2017).
18. Chmura, G. L., Anisfeld, S. C., Cahoon, D. R. & Lynch, J. C. Global carbon sequestration in tidal, saline wetland soils. *Glob. Biogeochem. Cycles* **17**, (2003).
19. Maxwell, T. L. *et al.* Global dataset of soil organic carbon in tidal marshes. *Sci. Data* **10**, 797 (2023).
20. Holmquist, J. R. *et al.* The Coastal Carbon Library and Atlas: Open source soil data and tools supporting blue carbon research and policy. *Glob. Change Biol.* **30**, e17098 (2024).
21. Mason, V. G. *et al.* Blue carbon benefits from global saltmarsh restoration. *Glob. Change Biol.* **29**, 6517–6545 (2023).
22. Duarte, C. M., Losada, I. J., Hendriks, I. E., Mazarrasa, I. & Marbà, N. The role of coastal

plant communities for climate change mitigation and adaptation. *Nat. Clim. Change* **3**, 961–968 (2013).
23. Smeaton, C. *et al.* Organic carbon stocks of Great British saltmarshes. *Front. Mar. Sci.* **10**, (2023).
24. Hansen, K. *et al.* Factors influencing the organic carbon pools in tidal marsh soils of the Elbe estuary (Germany). *J. Soils Sediments* **17**, 47–60 (2017).
25. Artigas, F. *et al.* Long term carbon storage potential and CO2 sink strength of a restored salt marsh in New Jersey. *Agric. For. Meteorol.* **200**, 313–321 (2015).
26. Alongi, D. M. Carbon balance in salt marsh and mangrove ecosystems: A global synthesis. *J. Mar. Sci. Eng.* **8**, 767 (2020).
27. Maxwell, T. L. *et al.* Global mangrove soil organic carbon stocks dataset at 30 m resolution for the year 2020 based on spatiotemporal predictive machine learning. *Data Brief* **50**, 109621 (2023).
28. Goldstein, A. *et al.* Protecting irrecoverable carbon in Earth's ecosystems. *Nat. Clim. Change* **10**, 287–295 (2020).
29. Serrano, O. *et al.* Australian vegetated coastal ecosystems as global hotspots for climate change mitigation. *Nat. Commun.* **10**, 4313 (2019).
30. Smith, A. J., Noyce, G. L., Megonigal, J. P., Guntenspergen, G. R. & Kirwan, M. L. Temperature optimum for marsh resilience and carbon accumulation revealed in a whole-ecosystem warming experiment. *Glob. Change Biol.* **28**, 3236–3245 (2022).
31. Rogers, K. *et al.* Wetland carbon storage controlled by millennial-scale variation in relative sea-level rise. *Nature* **567**, 91–95 (2019).
32. Adhikari, S. & Ivins, E. R. Climate-driven polar motion: 2003–2015. *Sci. Adv.* **2**, e1501693 (2016).
33. Spalding, M. D. *et al.* Marine Ecoregions of the World: A Bioregionalization of Coastal and Shelf Areas. *BioScience* **57**, 573–583 (2007).
34. Meyer, H., Reudenbach, C., Wöllauer, S. & Nauss, T. Importance of spatial predictor variable selection in machine learning applications – Moving from data reproduction to spatial prediction. *Ecol. Model.* **411**, 108815 (2019).
35. Sanderman, J. *et al.* A global map of mangrove forest soil carbon at 30 m spatial resolution. *Environ. Res. Lett.* **13**, (2018).
36. Connor, R. F., Chmura, G. L. & Beecher, C. B. Carbon accumulation in Bay of Fundy salt marshes: Implications for restoration of reclaimed marshes. *Glob. Biogeochem. Cycles* **15**, 943–954 (2001).
37. Rogers, K., Macreadie, P. I., Kelleway, J. J. & Saintilan, N. Blue carbon in coastal landscapes: a spatial framework for assessment of stocks and additionality. *Sustain. Sci.* **14**, 453–467 (2019).
38. Gore, C. *et al.* Saltmarsh blue carbon accumulation rates and their relationship with sea-level rise on a multi-decadal timescale in northern England. *Estuar. Coast. Shelf Sci.* **299**, 108665 (2024).
39. Clark, J. A., Farrell, W. E. & Peltier, W. R. Global Changes in Postglacial Sea Level: A Numerical Calculation1. *Quat. Res.* **9**, 265–287 (1978).
40. Sharma, R., Mishra, D. R., Levi, M. R. & Sutter, L. A. Remote Sensing of Surface and Subsurface Soil Organic Carbon in Tidal Wetlands: A Review and Ideas for Future Research. *Remote Sens.* **14**, 2940 (2022).
41. Wang, W. *et al.* Species-Specific Impacts of Invasive Plant Success on Vertical Profiles of Soil Carbon Accumulation and Nutrient Retention in the Minjiang River Tidal Estuarine Wetlands of China. *Soil Syst.* **2**, (2018).

42. Martinetto, P. *et al.* The blue carbon of southern southwest Atlantic salt marshes and their biotic and abiotic drivers. *Nat. Commun.* **14**, 8500 (2023).
43. Rovai, A. S. *et al.* Global controls on carbon storage in mangrove soils. *Nat. Clim. Change* **8**, 534+ (2018).
44. Rogers, K. Accommodation space as a framework for assessing the response of mangroves to relative sea-level rise. *Singap. J. Trop. Geogr.* **42**, 163–183 (2021).
45. Meyer, H. & Pebesma, E. Predicting into unknown space? Estimating the area of applicability of spatial prediction models. *Methods Ecol. Evol.* **12**, 1620–1633 (2021).
46. Viswanathan, C. *et al.* Salt marsh vegetation in India: Species composition, distribution, zonation pattern and conservation implications. *Estuar. Coast. Shelf Sci.* **242**, 106792 (2020).
47. Ruiz-Fernandez, A. C. *et al.* Carbon burial and storage in tropical salt marshes under the influence of sea level rise. *Sci. TOTAL Environ.* **630**, 1628–1640 (2018).
48. Perera, N., Lokupitiya, E., Halwatura, D. & Udagedara, S. Quantification of blue carbon in tropical salt marshes and their role in climate change mitigation. *Sci. Total Environ.* **820**, 153313 (2022).
49. McGuire, A. D. *et al.* Sensitivity of the carbon cycle in the Arctic to climate change. *Ecol. Monogr.* **79**, 523–555 (2009).
50. Chastain, S. G., Kohfeld, K. E., Pellatt, M. G., Olid, C. & Gailis, M. Quantification of blue carbon in salt marshes of the Pacific coast of Canada. *Biogeosciences* **19**, 5751–5777 (2022).
51. Maxwell, T. L. *et al.* Database: Tidal Marsh Soil Organic Carbon (MarSOC) Dataset. Zenodo https://doi.org/10.5281/zenodo.8414110 (2023).
52. Holmquist, J., Wolfe, J., Lonneman, M., Klinges, D. & Megonigal, J. P. Database: Coastal Carbon Network Data Library. (2022) doi:https://doi.org/10.25573/serc.21565671.
53. Murray, N. J. *et al.* High-resolution mapping of losses and gains of Earth's tidal wetlands. *Science* **376**, 744–749 (2022).
54. Abbott, K. M., Quirk, T. & Delaune, R. D. Dataset: Factors influencing blue carbon accumulation across a 32-year chronosequence of created coastal marshes. (2019) doi:10.25573/DATA.10005215.
55. Abbott, K. M., Elsey-Quirk, T. & DeLaune, R. D. Factors influencing blue carbon accumulation across a 32-year chronosequence of created coastal marshes. Ecosphere 10, (2019).
56. Adame, M. F. et al. Mangroves in arid regions: Ecology, threats, and opportunities. Estuarine, Coastal and Shelf Science 248, 106796 (2021).
57. Adame, M. F. et al. Carbon and Nitrogen Sequestration of Melaleuca Floodplain Wetlands in Tropical Australia. Ecosystems 23, 454–466 (2020).
58. Adame, M. F. et al. Carbon stocks and soil sequestration rates of tropical riverine wetlands. Biogeosciences 12, 3805–3818 (2015).
59. Adame, M. F. et al. Carbon Stocks of Tropical Coastal Wetlands within the Karstic Landscape of the Mexican Caribbean. PLOS ONE 8, e56569 (2013).
60. Allen, J. R., Cornwell, J. C. & Baldwin, A. H. Dataset: Contributions of organic and mineral matter to vertical accretion in tidal wetlands across a chesapeake bay subestuary. (2022) doi:10.25573/serc.18130892.v1.
61. Allen, J. R., Cornwell, J. C. & Baldwin, A. H. Contributions of organic and mineral matter to vertical accretion in tidal wetlands across a chesapeake bay subestuary. Journal of Marine Science and Engineering 9, (2021).

62. Anisfeld, S. C., Tobin, M. J. & Benoit, G. Sedimentation Rates in Flow-Restricted and Restored Salt Marshes in Long Island Sound. Estuaries 22, 231 (1999).
63. Arias-Ortiz, A., MasqueNANA, P., Paytan, A. & Baldocch, D. D. Dataset: Tidal and nontidal marsh restoration: a trade-off between carbon sequestration, methane emissions, and soil accretion. (2021) doi:10.25573/serc.15127743.v2.
64. Arias-Ortiz, A. et al. Tidal and nontidal marsh restoration: a trade-off between carbon sequestration, methane emissions, and soil accretion. Journal of Geophysical Research: Biogeosciences (2021) doi:10.1029/2021JG006573.
65. Arriola, J. M. & Cable, J. E. Variations in carbon burial and sediment accretion along a tidal creek in a Florida salt marsh. Limnology and Oceanography 62, S15–S28 (2017).
66. Baustian, M. M. et al. Long-term soil carbon data and accretion from four marsh types in Mississippi River Delta in 2015. (2021) doi:10.5066/P93U3B3E.
67. Baustian, M. M., Stagg, C. L., Perry, C. L., Moss, L. C. & Carruthers, T. J. B. Long-term carbon sinks in marsh soils of coastal louisiana are at risk to wetland loss. Journal of Geophysical Research: Biogeosciences 126, (2021).
68. Beasy, K. & Ellison, J. Comparison of Three Methods for the Quantification of Sediment Organic Carbon in Salt Marshes of the Rubicon Estuary, Tasmania, Australia. International Journal of Biology 5, p1 (2013).
69. Bernhardt, C. E. et al. Carbon budget assessment of tidal freshwater forested wetland and oligohaline marsh ecosystems along the Waccamaw and Savannah rivers, U.S.A. (2005-2016). (2018) doi:10.5066/F7TM7930.
70. Boyd, B. Comparison of sediment accumulation and accretion in impounded and unimpounded marshes of the Delaware Estuary. (University of Delaware, 2012).
71. Boyd, B. M. & Sommerfield, C. K. Marsh accretion and sediment accumulation in a managed tidal wetland complex of Delaware Bay. Ecological Engineering 92, 37–46 (2016).
72. Boyd, B. M., Sommerfield, C. K. & Elsey-Quirk, T. Hydrogeomorphic influences on salt marsh sediment accumulation and accretion in two estuaries of the U.S. Mid-Atlantic coast. Marine Geology 383, 132–145 (2017).
73. Boyd, B., Sommerfield, C. K., Quirk, T. & Unger, V. Dataset: Accretion and sediment accumulation in impounded and unimpounded marshes in the Delaware Estuary and Barnegat Bay. (2019) doi:10.25573/DATA.9747065.
74. Breithaupt et al. Dataset: Increasing rates of carbon burial in southwest Florida coastal wetlands. (2020) doi:10.25573/data.9894266.
75. Bryant, J. C. & Chabreck, R. H. Effects of Impoundment on Vertical Accretion of Coastal Marsh. Estuaries 21, 416 (1998).
76. Bulmer, R. H. et al. Blue Carbon Stocks and Cross-Habitat Subsidies. Frontiers in Marine Science 7, 380 (2020).
77. Bunzel, D. et al. (Table A1) Organic carbon measurements for sediment sequences TB13-1, GeoHH-GIE, GeoHH-FK and GeoHH-KWK. In supplement to: Bunzel, D et al. (2020): Integrated stratigraphy of foreland salt-marsh sediments of the south-eastern North Sea region. Newsletters on Stratigraphy, https://doi.org/10.1127/nos/2020/0540 PANGAEA https://doi.org/10.1594/PANGAEA.905218 (2019).
78. Burden, A., Garbutt, A. & Evans, C. D. Effect of restoration on saltmarsh carbon accumulation in Eastern England. Biology Letters 15, 20180773 (2019).
79. Burden, A., Garbutt, A., Hughes, S., Oakley, S. & Tempest, J. A. Soil biochemical measurements from salt marshes of different ages on the Essex coast, UK (2011). NERC Environmental Information Data Centre https://doi.org/10.5285/0b1faab4-3539-

457f-9169-b0b1fbd59bc2 (2018).
80. Burden, A., Garbutt, A., Hughes, S., Oakley, S. & Tempest, J. A. Soil biochemical measurements from salt marshes of different ages on the Essex coast, UK (2011). (2018) doi:10.5285/0B1FAAB4-3539-457F-9169-B0B1FBD59BC2.
81. Burke, S. A., Manahan, J., Eichelmann, E. & Cott, G. M. Dublin's saltmarshes contain climate-relevant carbon pools. Frontiers in Marine Science 9, 976457 (2022).
82. Cahoon, D. R., Lynch, J. C. & Powell, A. N. Marsh Vertical Accretion in a Southern California Estuary, U.S.A. Estuarine, Coastal and Shelf Science 43, 19–32 (1996).
83. Callaway, J. C., Borgnis, E. L., Turner, R. E. & Milan, C. S. Carbon sequestration and sediment accretion in san francisco bay tidal wetlands. Estuaries and Coasts 35, 1163–1181 (2012).
84. Callaway, J. C., Evyan L. Borgnis, R. Eugene Turner & Milan, C. S. Dataset: Carbon sequestration and sediment accretion in San Francisco Bay tidal wetlands. (2019) doi:10.25573/DATA.9693251.
85. Camacho, S., Moura, D., Connor, S., Boski, T. & Gomes, A. Geochemical characteristics of sediments along the margins of an atlantic-mediterranean estuary (the Guadiana, Southeast Portugal): spatial and seasonal variations. RGCI 14, 129–148 (2014).
86. Carlin, J. et al. Dataset: Sedimentary organic carbon measurements in a restored coastal wetland in san francisco bay, CA, USA. (2021) doi:10.25573/SERC.16416684.
87. Chambers, L. et al. Barataria Bay carbon mineralization and biogeochemical properties from nine soil cores. (2019) doi:10.1575/1912/bco-dmo.775547.1.
88. Chmura, G. L. & Hung, G. A. Controls on salt marsh accretion: A test in salt marshes of Eastern Canada. Estuaries 27, 70–81 (2004).
89. Cochran, J. K., Hirschberg, D. J., Wang, J. & Dere, C. Atmospheric deposition of metals to coastal waters (long island sound, new york U.S.A.): Evidence from saltmarsh deposits. Estuarine, Coastal and Shelf Science 46, 503–522 (1998).
90. Connor, R. F., Chmura, G. L. & Beecher, C. B. Carbon accumulation in bay of fundy salt marshes: Implications for restoration of reclaimed marshes. Global Biogeochem. Cycles 15, 943–954 (2001).
91. Conrad, S. et al. Does Regional Development Influence Sedimentary Blue Carbon Stocks? A Case Study From Three Australian Estuaries. Frontiers in Marine Science 5, 518 (2019).
92. Cott, G. M., Chapman, D. V. & Jansen, M. A. K. Salt Marshes on Substrate Enriched in Organic Matter: The Case of Ombrogenic Atlantic Salt Marshes. Estuaries and Coasts 36, 595–609 (2013).
93. Craft, C. B., Seneca, E. D. & Broome, S. W. Vertical Accretion in Microtidal Regularly and Irregularly Flooded Estuarine Marshes. Estuarine, Coastal and Shelf Science 37, 371–386 (1993).
94. Craft, C. Freshwater input structures soil properties, vertical accretion, and nutrient accumulation of Georgia and U.S tidal marshes. Limnology and Oceanography 52, 1220–1230 (2007).
95. Crooks, S. et al. Coastal Blue Carbon Opportunity Assessment for the Snohomish Estuary: The Climate Benefits of Estuary Restoration. doi.org/10.13140/RG.2.1.1371.6568 (2014) doi:10.13140/RG.2.1.1371.6568.
96. Cuellar-Martinez, T., Ruiz-Fernández, A. C., Sanchez-Cabeza, J.-A., Pérez-Bernal, L.-H. & Sandoval-Gil, J. Relevance of carbon burial and storage in two contrasting blue carbon ecosystems of a north-east Pacific coastal lagoon. Science of The Total Environment 675, 581–593 (2019).

97. Cuellar-Martinez, T. et al. Temporal records of organic carbon stocks and burial rates in Mexican blue carbon coastal ecosystems throughout the Anthropocene. Global and Planetary Change 192, 103215 (2020).
98. Cusack, M. et al. Organic carbon sequestration and storage in vegetated coastal habitats along the western coast of the Arabian Gulf. Environmental Research Letters 13, 074007 (2018).
99. Day, J. W. et al. Vegetation death and rapid loss of surface elevation in two contrasting Mississippi delta salt marshes: The role of sedimentation, autocompaction and sea-level rise. Ecological Engineering 37, 229–240 (2011).
100. de los Santos, C. B. et al. Sedimentary Organic Carbon and Nitrogen Sequestration Across a Vertical Gradient on a Temperate Wetland Seascape Including Salt Marshes, Seagrass Meadows and Rhizophytic Macroalgae Beds. Ecosystems 26, 826–842 (2023).
101. de los Santos, C. B. et al. Vertical intertidal variation of organic matter stocks and patterns of sediment deposition in a mesotidal coastal wetland. Estuarine, Coastal and Shelf Science 272, 107896 (2022).
102. Doughty, C. et al. Mangroves marching northward: the impacts of rising seas and temperatures on ecosystems at Kennedy Space Center. (2019) doi:10.25573/DATA.9695918.V1.
103. Doughty, C. L. et al. Mangrove range expansion rapidly increases coastal wetland carbon storage. Estuaries and Coasts 39, 385–396 (2015).
104. Drake, K., Halifax, H., Adamowicz, S. C. & Craft, C. Carbon sequestration in tidal salt marshes of the northeast united states. Environmental Management 56, 998–1008 (2015).
105. Drexler, J. Z. et al. A long-term comparison of carbon sequestration rates in impounded and naturally tidal freshwater marshes along the lower waccamaw river, south carolina. Wetlands 33, 965–974 (2013).
106. Drexler, J. Z., Woo, I., Fuller, C. C. & Nakai, G. Carbon accumulation and vertical accretion in a restored versus historic salt marsh in southern Puget Sound, Washington, United States. Restoration Ecology 27, 1117–1127 (2019).
107. Elsey-Quirk, T., Seliskar, D. M., Sommerfield, C. K. & Gallagher, J. L. Salt marsh carbon pool distribution in a mid-atlantic lagoon, USA: Sea level rise implications. Wetlands 31, 87–99 (2011).
108. Ensign, S. H., Noe, G. B., Hupp, C. R. & Skalak, K. J. Head-of-tide bottleneck of particulate material transport from watersheds to estuaries. Geophysical Research Letters 42, 10,671-10,679 (2015).
109. Ensign, S. H., Noe, G. B., Hupp, C. R. & Skalak, K. J. Dataset: Head of tide bottleneck of particulate material transport from watersheds to estuaries. (2021) doi:10.25573/SERC.13483332.
110. Ewers Lewis, C. J., Carnell, P. E., Sanderman, J., Baldock, J. A. & Macreadie, P. I. Variability and Vulnerability of Coastal 'Blue Carbon' Stocks: A Case Study from Southeast Australia. Ecosystems 21, 263–279 (2018).
111. Fell, C., Adgie, T. & Chapman, S. Dataset: Carbon sequestration across northeastern, florida coastal wetlands. (2021) doi:10.25573/serc.15043920.v1.
112. Ferronato, C. et al. Effect of waterlogging on soil biochemical properties and organic matter quality in different salt marsh systems. Geoderma 338, 302–312 (2019).
113. Ford, H., Garbutt, A. & Skov, M. Coastal Biodiversity and Ecosystem Service Sustainability (CBESS) soil organic matter content from three soil depths on saltmarsh

sites at Morecambe Bay and Essex. NERC Environmental Information Data Centre https://doi.org/10.5285/90457ba1-f291-4158-82dc-425d7cbb1ac5 (2016).
114. Fu, C. et al. Stocks and losses of soil organic carbon from Chinese vegetated coastal habitats. Global Change Biology 27, 202–214 (2021).
115. Fuchs, M. et al. Soil carbon and nitrogen stocks in Arctic river deltas: New data for three Northwest Alaskan deltas. in (Chamonix Mont-Blanc, France, 2018).
116. Gailis, M., Kohfeld, K. E., Pellatt, M. G. & Carlson, D. Quantifying blue carbon for the largest salt marsh in southern British Columbia: implications for regional coastal management. Coastal Engineering Journal 63, 275–309 (2021).
117. Gallagher, J. B., Prahalad, V. & Aalders, J. Inorganic and Black Carbon Hotspots Constrain Blue Carbon Mitigation Services Across Tropical Seagrass and Temperate Tidal Marshes. Wetlands 41, 65 (2021).
118. Gerlach, M. J. et al. Reconstructing Common Era relative sea-level change on the Gulf Coast of Florida. Marine Geology 390, 254–269 (2017).
119. Giblin, A., Forbrich, I., & Plum Island Ecosystems LTER. PIE LTER high marsh sediment chemistry and activity measurements, Nelson Island Creek marsh, Rowley, MA. (2018) doi:10.6073/PASTA/D1D5CBF87602CCF51DE30B87B8E46D01.
120. Gispert, M. et al. Appraising soil carbon storage potential under perennial and annual Chenopodiaceae in salt marsh of NE Spain. Estuarine, Coastal and Shelf Science 252, 107240 (2021).
121. Gispert, M., Phang, C. & Carrasco-Barea, L. The role of soil as a carbon sink in coastal salt-marsh and agropastoral systems at La Pletera, NE Spain. CATENA 185, 104331 (2020).
122. Gonneea, M. E., Kroeger, K. D., & and O&Amp. Collection, analysis, and age-dating of sediment cores from salt marshes on the south shore of Cape Cod, Massachusetts, from 2013 through 2014. (2018) doi:10.5066/F7H41QPP.
123. González-Alcaraz, M. N., Aránega, B., Conesa, H. M., Delgado, M. J. & Álvarez-Rogel, J. Contribution of soil properties to the assessment of a seawater irrigation programme as a management strategy for abandoned solar saltworks. CATENA 126, 189–200 (2015).
124. González-Alcaraz, M. N. et al. Storage of organic carbon, nitrogen and phosphorus in the soil–plant system of Phragmites australis stands from a eutrophicated Mediterranean salt marsh. Geoderma 185–186, 61–72 (2012).
125. Gorham, C., Lavery, P., Kelleway, J. J., Salinas, C. & Serrano, O. Soil Carbon Stocks Vary Across Geomorphic Settings in Australian Temperate Tidal Marsh Ecosystems. Ecosystems 24, 319–334 (2021).
126. Graversen, A. E. L., Banta, G. T., Masque, P. & Krause-Jensen, D. Carbon sequestration is not inhibited by livestock grazing in Danish salt marshes. Limnology and Oceanography 67, S19–S35 (2022).
127. Grey, A. et al. Geochemical mapping of a blue carbon zone: Investigation of the influence of riverine input on tidal affected zones in Bull Island. Regional Studies in Marine Science 45, 101834 (2021).
128. Gu, J., vanArdenne, L. & Chmura, G. Data for: Invasive Phragmites increases blue carbon stock and soil volume in a St. Lawrence estuary marsh. Mendeley Data (2020).
129. Guerra, R., Simoncelli, S. & Pasteris, A. Carbon accumulation and storage in a temperate coastal lagoon under the influence of recent climate change (Northwestern Adriatic Sea). Regional Studies in Marine Science 53, 102439 (2022).
130. Hansen, K. et al. Factors influencing the organic carbon pools in tidal marsh soils of the

Elbe estuary (Germany). Journal of Soils and Sediments 17, 47–60 (2017).
131. Hatje, V. et al. Vegetated coastal ecosystems in the Southwestern Atlantic Ocean are an unexploited opportunity for climate change mitigation. Communications Earth & Environment 4, 1–10 (2023).
132. Hatton, R. S., DeLaune, R. D. & Patrick, W. H. Sedimentation, accretion, and subsidence in marshes of Barataria Basin, Louisiana1: Marsh accretion, subsidence. Limnology and Oceanography 28, 494–502 (1983).
133. Hayes, M. A. et al. Dynamics of sediment carbon stocks across intertidal wetland habitats of Moreton Bay, Australia. Global Change Biology 23, 4222–4234 (2017).
134. He, Q. et al. Consumer regulation of the carbon cycle in coastal wetland ecosystems. Philosophical Transactions of the Royal Society B: Biological Sciences 375, 20190451 (2020).
135. Hill, T. D. & Anisfeld, S. C. Coastal wetland response to sea level rise in Connecticut and New York. Estuarine, Coastal and Shelf Science 163, 185–193 (2015).
136. Holmquist, J. R. et al. Accuracy and precision of tidal wetland soil carbon mapping in the conterminous united states: Public soil carbon data release. (2018) doi:10.25572/ccrcn/10088/35684.
137. Hu, M., Sardans, J., Yang, X., Peñuelas, J. & Tong, C. Patterns and environmental drivers of greenhouse gas fluxes in the coastal wetlands of China: A systematic review and synthesis. Environmental Research 186, 109576 (2020).
138. Human, L. R. D., Els, J., Wasserman, J. & Adams, J. B. Blue carbon and nutrient stocks in salt marsh and seagrass from an urban African estuary. Science of The Total Environment 842, 156955 (2022).
139. J. Boone Kauffman et al. Dataset: Carbon stocks in seagrass meadows, emergent marshes, and forested tidal swamps of the Pacific Northwest. (2020) doi:10.25573/serc.12640172.
140. Johnson, B. J., Moore, K. A., Lehmann, C., Bohlen, C. & Brown, T. A. Middle to late holocene fluctuations of C3 and C4 vegetation in a northern new england salt marsh, sprague marsh, phippsburg maine. Organic Geochemistry 38, 394–403 (2007).
141. Jones, M. C., Bernhardt, C. E., Krauss, K. W. & Noe, G. B. The impact of late holocene land use change, climate variability, and sea level rise on carbon storage in tidal freshwater wetlands on the southeastern united states coastal plain. Journal of Geophysical Research: Biogeosciences 122, 3126–3141 (2017).
142. Karen Thorne, U. S. G. S. Marshes to mudflats: Climate change effects along a latitudinal gradient in the pacific northwest. (2015) doi:10.5066/F7SJ1HNC.
143. Kauffman, J. B. et al. Total ecosystem carbon stocks at the marine-terrestrial interface: Blue carbon of the Pacific Northwest Coast, United States. Global Change Biology (2020) doi:10.1111/gcb.15248.
144. Kauffman, J. B. et al. SWAMP Dataset-Mangrove soil carbon-marisma-2017. CIFOR https://doi.org/10.17528/CIFOR/DATA.00244 (2020).
145. Kemp, A. C. et al. Use of lead isotopes for developing chronologies in recent salt-marsh sediments. Quaternary Geochronology 12, 40–49 (2012).
146. Kemp, A. C. et al. Dataset: Use of lead isotopes for developing chronologies in recent salt-marsh sediments. (2020) doi:10.25573/serc.11569419.
147. Keshta, A. E., Yarwood, S. A. & Baldwin, A. H. Hydrology, soil redox, and pore-water iron regulate carbon cycling in natural and restored tidal freshwater wetlands in the chesapeake bay, maryland, usa. (University of Maryland, 2017). doi:10.13016/M2S756M71.

148. Keshta, A. E., Yarwood, S. A. & Baldwin, A. H. Dataset: Soil redox and hydropattern control soil carbon stocks across different habitats in tidal freshwater wetlands in a sub-estuary of the chesapeake bay. (2020) doi:10.25573/serc.13187549.
149. Keshta, A. E., Yarwood, S. A. & Baldwin, A. H. A new in situ method showed greater persistence of added soil organic matter in natural than restored wetlands. Restoration Ecology (2021) doi:10.1111/rec.13437.
150. Kohfeld, K. E., Chastain, S., Pellatt, M. G. & Olid, C. Salt marsh soil carbon content, loss on ignition, dry bulk density, carbon stocks and carbon accumulation rates for Clayoquot Sound, British Columbia, Canada. In: Kohfeld, KE et al. (2022): Salt marsh soil carbon content, loss on ignition, dry bulk density, carbon stocks, lead-210 and carbon accumulation rates, for Clayoquot Sound, British Columbia, Canada. PANGAEA, https://doi.org/10.1594/PANGAEA.947824 (2022).
151. Krauss, K. W. et al. The role of the upper tidal estuary in wetland blue carbon storage and flux. Global Biogeochemical Cycles 32, 817–839 (2018).
152. Kulawardhana, R. W. et al. The role of elevation, relative sea-level history and vegetation transition in determining carbon distribution in Spartina alterniflora dominated salt marshes. Estuarine, Coastal and Shelf Science 154, 48–57 (2015).
153. Kumar, M., Boski, T., González-Vila, F. J., Jiménez-Morillo, N. T. & González-Pérez, J. A. Characteristics of organic matter sources from Guadiana Estuary salt marsh sediments (SW Iberian Peninsula). Continental Shelf Research 197, 104076 (2020).
154. Lagomasino, D., Corbett, D. R. & Walsh, J. Influence of wind-driven inundation and coastal geomorphology on sedimentation in two microtidal marshes, Pamlico River Estuary, NC. Estuaries and coasts 36, 1165–1180 (2013).
155. Lagomasino, D., D. Reide Corbett, & J.P. Walsh. Dataset: Influence of wind-driven inundation and coastal geomorphology on sedimentation in two microtidal marshes, pamlico river estuary, NC. (2020) doi:10.25573/SERC.12043335.
156. Laurent, K. A. St., Hribar, D. J., Carlson, A. J., Crawford, C. M. & Siok, D. Assessing coastal carbon variability in two Delaware tidal marshes. Journal of Coastal Conservation 24, (2020).
157. Laurent, K. A. St., Hribar, D. J., Carlson, A. J., Calyn M. Crawford, & Drexel Siok. Dataset: Assessing coastal carbon variability in two Delaware tidal marshes. (2020) doi:10.25573/SERC.13315472.
158. Li, Y. et al. Plant biomass and soil organic carbon are main factors influencing dry-season ecosystem carbon rates in the coastal zone of the Yellow River Delta. PLOS ONE 14, e0210768 (2019).
159. Loomis, M. J. & Craft, C. B. Carbon Sequestration and Nutrient (Nitrogen, Phosphorus) Accumulation in River-Dominated Tidal Marshes, Georgia, USA. Soil Science Society of America Journal 74, 1028–1036 (2010).
160. Luk, S., Spivak, A., Eagle, M. J. & O'keefe Suttles, J. A. Collection, analysis, and age-dating of sediment cores from a salt marsh platform and ponds, Rowley, Massachusetts, 2014-15. (2020) doi:10.5066/P9HIOWKT.
161. Luk, S. Y. et al. Soil organic carbon development and turnover in natural and disturbed salt marsh environments. Geophysical Research Letters 48, (2021).
162. Macreadie, P. I. et al. Carbon sequestration by Australian tidal marshes. Scientific Reports 7, 44071 (2017).
163. Markewich, H. W. et al. Detailed Descriptions for Sampling, Sample Preparation and Analyses of Cores from St. Bernard Parish, Louisiana. Open-File Report https://pubs.er.usgs.gov/publication/ofr98429 (1998) doi:10.3133/ofr98429.

164. Martins, M. et al. Carbon and Nitrogen Stocks and Burial Rates in Intertidal Vegetated Habitats of a Mesotidal Coastal Lagoon. Ecosystems 25, 372–386 (2022).
165. Mazarrasa, I. et al. Drivers of variability in Blue Carbon stocks and burial rates across European estuarine habitats. Science of The Total Environment 886, 163957 (2023).
166. McClellan, S. A. Data for: Root-zone carbon and nitrogen pools across two chronosequences of coastal marshes formed using different restoration techniques: Dredge sediment versus river sediment diversion. (2021) doi:10.17632/5ZBV2MB5ZP.1.
167. McClellan, S. A., Elsey-Quirk, T., Laws, E. A. & DeLaune, R. D. Root-zone carbon and nitrogen pools across two chronosequences of coastal marshes formed using different restoration techniques: Dredge sediment versus river sediment diversion. Ecological Engineering 169, 106326 (2021).
168. McTigue, N. et al. Dataset: Carbon accumulation rates in a salt marsh over the past two millennia. (2020) doi:10.25573/serc.11421063.
169. McTigue, N. et al. Sea-level rise explains changing carbon accumulation rates in a salt marsh over the past two millennia. Journal of Geophysical Research: Biogeosciences (2019) doi:10.1029/2019JG005207.
170. Merrill, J. Z. Tidal freshwater marshes as nutrient sinks: Particulate nutrient burial and denitrification. (University of Maryland, College Park, 1999).
171. Messerschmidt, T. C. & Kirwan, M. L. Dataset: Soil properties and accretion rates of C3 and C4 marshes at the global change research wetland, edgewater, maryland. (2020) doi:10.25573/SERC.11914140.
172. Miller, C. B., Rodriguez, A. B., Bost, M. C., McKee, B. A. & McTigue, N. D. Carbon accumulation rates are highest at young and expanding salt marsh edges. Communications Earth & Environment (2022) doi:10.1038/s43247-022-00501-x.
173. Miller, L. C., Smeaton, C., Yang, H. & Austin, W. E. N. Physical and geochemical properties of Scottish saltmarsh soils. Marine Scotland https://doi.org/10.7489/12422-1 (2022).
174. Morris, J. T. & Jensen, A. The carbon balance of grazed and non-grazed Spartina anglica saltmarshes at Skallingen, Denmark. Journal of Ecology 86, 229–242 (1998).
175. Nahlik, M., A., Fennessy, & Siobhan. Carbon storage in US wetlands. Nature Communications (2016) doi:doi.org/10.1038/ncomms13835.
176. Neubauer, S. C., Anderson, I. C., Constantine, J. A. & Kuehl, S. A. Sediment deposition and accretion in a mid-atlantic (U.S.A.) tidal freshwater marsh. Estuarine, Coastal and Shelf Science 54, 713–727 (2002).
177. Noe, G. B., Hupp, C. R., Bernhardt, C. E. & Krauss, K. W. Contemporary deposition and long-term accumulation of sediment and nutrients by tidal freshwater forested wetlands impacted by sea level rise. Estuaries and Coasts 39, 1006–1019 (2016).
178. Nogueira, J. et al. Geochemistry of coastal wetland in the Northern Saharan environment through lacustrine sediment core THI. PANGAEA PANGAEA https://doi.org/10.1594/PANGAEA.925024 (2020).
179. Nolte, S. Dataset: Does livestock grazing affect sediment deposition and accretion rates in salt marshes? (2020) doi:10.25573/SERC.11958996.
180. Nolte, S. et al. Does livestock grazing affect sediment deposition and accretion rates in salt marshes? Estuarine, Coastal and Shelf Science 135, 296–305 (2013).
181. Nuttle, W. Tidal Freshwater Marshes as Nutrient Sinks: Particulate Nutrient Burial and Denitrification. (Environmental Data Initiative, 1996).
182. Nyman, J., DeLaune, R., Roberts, H. & Patrick, W. Relationship between vegetation and soil formation in a rapidly submerging coastal marsh. Marine Ecology Progress

Series 96, 269–279 (1993).
183. O’keefe Suttles, J. A. et al. Collection, analysis, and age-dating of sediment cores from Herring River wetlands and other nearby wetlands in Wellfleet, Massachusetts, 2015-17. (2021) doi:10.5066/P95RXPHB.
184. O’keefe Suttles, J. A., Eagle, M. J., Mann, A. C. & Kroeger, K. D. Collection, analysis, and age-dating of sediment cores from mangrove and salt marsh ecosystems in Tampa Bay, Florida, 2015. (2021) doi:10.5066/P9QB17H2.
185. O’keefe Suttles, J. A. et al. Collection, analysis, and age-dating of sediment cores from natural and restored salt marshes on Cape Cod, Massachusetts, 2015-16. (2021) doi:10.5066/P9R154DY.
186. O’keefe Suttles, J. A. et al. Collection, analysis, and age-dating of sediment cores from salt marshes, rhode island, 2016. (2021) doi:10.5066/P94HIDVU.
187. Orson, R. A., Warren, R. S. & Niering, W. A. Interpreting Sea Level Rise and Rates of Vertical Marsh Accretion in a Southern New England Tidal Salt Marsh. Estuarine, Coastal and Shelf Science 47, 419–429 (1998).
188. Osland, M. J. Vegetation, soil, and landscape data. (2017) doi:10.5066/F7J1017G.
189. Pagès, J. F. et al. Resilience of saltmarsh carbon sequestration to ecosystem transitions. (in prep).
190. Pastore, M. A., Megonigal, J. P. & Langley, J. A. Elevated CO2 and nitrogen addition accelerate net carbon gain in a brackish marsh. Biogeochemistry 133, 73–87 (2017).
191. Patrick, Wm. H. & DeLaune, R. D. Subsidence. accretion. and sea level rise in south San Francisco Bay marshes. Limnol. Oceanogr. 35, 1389–1395 (1990).
192. Peck, E. K., Wheatcroft, R. A. & Brophy, L. S. Controls on sediment accretion and blue carbon burial in tidal saline wetlands: Insights from the oregon coast, USA. Journal of Geophysical Research: Biogeosciences 125, (2020).
193. Peck, E., Wheatcroft, R. & Brophy, L. Dataset: Controls on sediment accretion and blue carbon burial in tidal saline wetlands: Insights from the Oregon coast, U.S.A. (2020) doi:10.25573/SERC.11317820.V2.
194. Perera, N., Lokupitiya, E., Halwatura, D. & Udagedara, S. Quantification of blue carbon in tropical salt marshes and their role in climate change mitigation. Science of The Total Environment 820, 153313 (2022).
195. Piazza, S. C. et al. Geomorphic and ecological effects of Hurricanes Katrina and Rita on coastal Louisiana marsh communities. i–126 (2011) doi:10.3133/ofr20111094.
196. Pollmann, T., Böttcher, M. E. & Giani, L. Young soils of a temperate barrier island under the impact of formation and resetting by tides and wind. CATENA 202, 105275 (2021).
197. Poppe, K. L. & Rybczyk, J. M. Dataset: Sediment carbon stocks and sequestration rates in the Pacific Northwest region of Washington, USA. (2019) doi:10.25573/DATA.10005248.
198. Poppe, K. & Rybczyk, J. Tidal marsh restoration enhances sediment accretion and carbon accumulation in the Stillaguamish River estuary, Washington. Plos One (2021) doi:10.1371/journal.pone.0257244.
199. Radabaugh, K. R. et al. Coastal blue carbon assessment of mangroves, salt marshes, and salt barrens in tampa bay, florida, USA. Estuaries and Coasts 41, 1496–1510 (2017).
200. Rathore, A. P., Chaudhary, D. R. & Jha, B. Biomass production, nutrient cycling, and carbon fixation by Salicornia brachiata Roxb.: A promising halophyte for coastal saline soil rehabilitation. International Journal of Phytoremediation 18, 801–811 (2016).
201. Raw, J. et al. Salt marsh elevation and responses to future sea-level rise in the Knysna

Estuary, South Africa. African Journal of Aquatic Science 45, 49–64 (2020).
202. Richard A. Orson, R. L. S. Rates of sediment accumulation in a tidal freshwater marsh. SEPM Journal of Sedimentary Research (1990) doi:10.1306/d4267631-2b26-11d7-8648000102c1865d.
203. Rodriguez, Miller, A., Bost, C., & Molly. Salt marsh radiocarbon and loss on Ignition data. (2022) doi:10.6084/m9.figshare.20137649.v2.
204. Roman, C. T., Peck, J. A., Allen, J. R., King, J. W. & Appleby, P. G. Accretion of a New England (U.S.A.) Salt Marsh in Response to Inlet Migration, Storms, and Sea-level Rise. Estuarine, Coastal and Shelf Science 45, 717–727 (1997).
205. Ruranska, P., Ladd, C. J. T., Smeaton, C., Skov, M. W. & Austin, W. E. N. Dry bulk density, loss on ignition and organic carbon content of surficial soils from English and Welsh salt marshes 2019. NERC Environmental Information Data Centre https://doi.org/10.5285/e5554b83-910f-4030-8f4e-81967dc7047c (2022).
206. Ruranska, P. et al. Dry bulk density, loss on ignition and organic carbon content of surficial soils from Scottish salt marshes, 2018-2019. NERC Environmental Information Data Centre https://doi.org/10.5285/81a1301f-e5e2-44f9-afe0-0ea5bb08010f (2020).
207. Russell, S. K., Gillanders, B. M., Detmar, S., Fotheringham, D. & Jones, A. R. Determining Environmental Drivers of Fine-Scale Variability in Blue Carbon Soil Stocks. Estuaries and Coasts (2023) doi:10.1007/s12237-023-01260-4.
208. Rybczyk, J. M. & Cahoon, D. R. Estimating the potential for submergence for two wetlands in the Mississippi River Delta. Estuaries 25, 985–998 (2002).
209. Sammul, M., Kauer, K. & Köster, T. Biomass accumulation during reed encroachment reduces efficiency of restoration of Baltic coastal grasslands. Applied Vegetation Science 15, 219–230 (2012).
210. Sanbor, Paul, Coxson, & Darwyn. Dataset: Carbon data for intertidal soils and sediments, Skeena River estuary, British Columbia. (2020) doi:10.25573/serc.12252005.
211. Santos, R. et al. Superficial sedimentary stocks and sources of carbon and nitrogen in coastal vegetated assemblages along a flow gradient. Scientific Reports 9, 610 (2019).
212. Schile, L., Kauffman, J. B., Megonigal, J. P., Fourqurean, J. & Crooks, S. Abu Dhabi Blue Carbon project. Dryad https://doi.org/10.15146/R3K59Z (2016).
213. Schile, L. M. & Megonigal, J. P. Abu dhabi blue carbon demonstration project. (2017) doi:10.5479/data_serc/10088/31949.
214. Serrano, O. Soil carbon cores from Australian saltmarshes. (unpublished).
215. Serrano, O. et al. Australian vegetated coastal ecosystems as global hotspots for climate change mitigation. Nature Communications 10, 4313 (2019).
216. Shamrikova, E. V., Deneva, S. V. & Kubik, O. S. Spatial Patterns of Carbon and Nitrogen in Soils of the Barents Sea Coastal Area (Khaypudyrskaya Bay). Eurasian Soil Science 52, 507–517 (2019).
217. Siewert, M. B., Hugelius, G., Heim, B. & Faucherre, S. Landscape controls and vertical variability of soil organic carbon storage in permafrost-affected soils of the Lena River Delta. CATENA 147, 725–741 (2016).
218. Smeaton, C. et al. Physical and geochemical properties of saltmarsh soils from narrow diameter gouge cores in UK saltmarshes collected between 2018 and 2021. NERC EDS Environmental Information Data Centre https://doi.org/10.5285/d301c5f5-77f5-41ba-934e-a80e1293d4cd (2022).
219. Smeaton, C. et al. Physical and geochemical properties of saltmarsh soils from wide diameter gouge cores in UK saltmarshes collected between 2018 and 2021. NERC EDS Environmental Information Data Centre https://doi.org/10.5285/279558cd-20fb-4f19-

8077-4400817a4482 (2022).
220. Smeaton, C. et al. Physical and geochemical properties of saltmarsh soils from wide diameter gouge cores in Essex, UK, collected in 2019. NERC EDS Environmental Information Data Centre https://doi.org/10.5285/fa3f4087-528e-4c5d-90d8-6bb4675d6317 (2023).
221. Smeaton, C., Rees-Hughes, L., Barlow, N. L. M. & Austin, W. E. N. Sedimentological and organic carbon data from the Kyle of Tongue saltmarsh, Scotland, 2018. NERC Environmental Information Data Centre https://doi.org/10.5285/b57ef444-54d4-47f9-8cbf-3cfef1182b55 (2021).
222. Smith, K. E. L., Flocks, J. G., Steyer, G. D. & Piazza, S. C. Wetland paleoecological study of southwest coastal Louisiana: sediment cores and diatom calibration dataset. (2015) doi:10.3133/ds877.
223. Smith, K. E. L. Paleoecological study of coastal marsh in the Chenier Plain, Louisiana: investigating the diatom composition of hurricane-deposited sediments and a diatom-based quantitative reconstruction of sea-level characteristics. (University of Florida, 2012).
224. Sousa, A. I., Lillebø, A. I., Pardal, M. A. & Caçador, I. The influence of Spartina maritima on carbon retention capacity in salt marshes from warm-temperate estuaries. Marine Pollution Bulletin 61, 215–223 (2010).
225. Spivak, A. Bulk soil and elemental properties of marsh and infilled pond soils collected in 2014-2015 within Plum Island Ecosystems LTER. (2020) doi:10.26008/1912/BCO-DMO.827298.1.
226. Sun, H. et al. Soil organic carbon stabilization mechanisms in a subtropical mangrove and salt marsh ecosystems. Science of The Total Environment 673, 502–510 (2019).
227. Thom, R. M. Accretion rates of low intertidal salt marshes in the Pacific Northwest. Wetlands 12, 147–156 (1992).
228. Thom, R. M. Dataset: Accretion rates of low intertidal salt marshes in the Pacific Northwest. (2019) doi:10.25573/DATA.10046189.
229. Thorne, K. et al. U.S. Pacific coastal wetland resilience and vulnerability to sea-level rise. Science Advances 4, eaao3270 (2018).
230. Unger, V., Elsey-Quirk, T., Sommerfield, C. & Velinsky, D. Stability of organic carbon accumulating in Spartina alterniflora-dominated salt marshes of the Mid-Atlantic U.S. Estuarine, Coastal and Shelf Science 182, 179–189 (2016).
231. Van de Broek, M. Van de Broek et al. 2018, GCB, Supplementary data. Mendeley Data https://doi.org/10.17632/2nnv9bw3hh.2 (2018).
232. Vaughn, D., Bianchi, T., Shields, M., Kenney, W. & Osborne, T. Dataset: Increased organic carbon burial in northern florida mangrove-salt marsh transition zones. (2021) doi:10.25573/serc.10552004.v2.
233. Vaughn, D. R., Bianchi, T. S., Shields, M. R., Kenney, W. F. & Osborne, T. Z. Increased organic carbon burial in northern florida mangrove-salt marsh transition zones. Global Biogeochemical Cycles (2020) doi:10.1029/2019GB006334.
234. Vitti, S., Pellegrini, E., Casolo, V., Trotta, G. & Boscutti, F. Contrasting responses of native and alien plant species to soil properties shed new light on the invasion of dune systems. Journal of Plant Ecology 13, 667–675 (2020).
235. Voltz, B. et al. A multiproxy study of intertidal surface sediments from two macrotidal estuarine systems (Canche, Authie) in northern France: Insights into environmental processes. Continental Shelf Research 230, 104554 (2021).
236. Wails, C. N. et al. Assessing changes to ecosystem structure and function following

invasion by Spartina alterniflora and Phragmites australis: a meta-analysis. Biological Invasions 23, 2695–2709 (2021).
237. Ward, M. A. et al. Blue carbon stocks and exchanges along the California coast. Biogeosciences 18, 4717–4732 (2021).
238. Ward, M. Data from: Organic carbon, grain size, elemental/isotopic composition. (2021) doi:10.5061/dryad.m0cfxpp31.
239. Ward, M. A. et al. Blue carbon stocks and exchanges along the California coast. Biogeosciences 18, 4717–4732 (2021).
240. Ward, R. D. Carbon sequestration and storage in Norwegian Arctic coastal wetlands: Impacts of climate change. Science of The Total Environment 748, 141343 (2020).
241. Watson, E. B. & Byrne, R. Late holocene marsh expansion in southern san francisco bay, california. Estuaries and Coasts 36, 643–653 (2013).
242. Weis & Anthony, D. Vertical accretion rates and heavy metal chronologies in wetland sediments of Tijuana Estuary. (San Diego State University, 1999).
243. Weis, D. A., Callaway, J. C. & Gersberg, R. M. Vertical accretion rates and heavy metal chronologies in wetland sediments of the tijuana estuary. Estuaries 24, 840 (2001).
244. Weston, N. B. et al. Dataset: Recent acceleration of coastal wetland accretion along the U.S. east coast. (2022) doi:10.25573/serc.13043054.v1.
245. White, J. R., Sapkota, Y., Chambers, L. G., Cook, R. L. & Xue, Z. Biogeochemical properties of sediment cores from Barataria Basin, Louisiana, 2018 and 2019. (2020) doi:10.26008/1912/BCO-DMO.833824.1.
246. Windham-Myers, L. et al. Biogeochemical processes in an urban, restored wetland of San Francisco Bay, California, 2007-2009; methods and data for plant, sediment and water parameters. (2010) doi:10.3133/ofr20101299.
247. Wollenberg, J. T., Ollerhead, J. & Chmura, G. L. Rapid carbon accumulation following managed realignment on the Bay of Fundy. PLOS ONE 13, e0193930 (2018).
248. Xia, S. et al. Storage, patterns and influencing factors for soil organic carbon in coastal wetlands of China. Global Change Biology 28, 6065–6085 (2022).
249. Yando, E. S. et al. Salt marsh-mangrove ecotones: using structural gradients to investigate the effects of woody plant encroachment on plant–soil interactions and ecosystem carbon pools. Journal of Ecology 104, 1020–1031 (2016).
250. Yang, R.-M. & Chen, L.-M. Spartina alterniflora invasion alters soil bulk density in coastal wetlands of China. Land Degradation & Development 32, 1993–1999 (2021).
251. Ye, S. et al. Carbon Sequestration and Soil Accretion in Coastal Wetland Communities of the Yellow River Delta and Liaohe Delta, China. Estuaries and Coasts 38, 1885–1897 (2015).
252. Yu, O. T. & Chmura, G. L. Soil carbon may be maintained under grazing in a St Lawrence Estuary tidal marsh. Environmental Conservation 36, 312–320 (2009).
253. Yuan, H.-W. et al. Sources and distribution of sedimentary organic matter along the Andong salt marsh, Hangzhou Bay. Journal of Marine Systems 174, 78–88 (2017).
254. Yuan, J. et al. Data from: Spartina alterniflora invasion drastically increases methane production potential by shifting methanogenesis from hydrogenotrophic to methylotrophic pathway in a coastal marsh. Dryad https://doi.org/10.5061/DRYAD.6F60V3Q (2019).
255. Zubrzycki, S., Kutzbach, L., Grosse, G., Desyatkin, A. & Pfeiffer, E.-M. Organic carbon and total nitrogen stocks in soils of the Lena River Delta. Biogeosciences 10, 3507–3524 (2013).
256. National wetland condition assessment 2011. (2016).

257. Hengl, T. & MacMillan, R. A. *Predictive Soil Mapping with R*. (OpenGeoHub foundation, Wageningen, the Netherlands, 2019).
258. Murray, N. nick-murray/gee-global-intertidal-change-ARCHIVE: v1.0 for manuscript publication. Zenodo https://doi.org/10.5281/zenodo.6503069 (2022).
259. Gorelick, N. *et al.* Google Earth Engine: Planetary-scale geospatial analysis for everyone. *Remote Sens. Environ.* (2017) doi:10.1016/j.rse.2017.06.031.
260. Allen, J. R. L. Morphodynamics of Holocene salt marshes: a review sketch from the Atlantic and Southern North Sea coasts of Europe. *Quat. Sci. Rev.* **19**, 1155–1231 (2000).
261. Spivak, A. C., Sanderman, J., Bowen, J. L., Canuel, E. A. & Hopkinson, C. S. Global-change controls on soil-carbon accumulation and loss in coastal vegetated ecosystems. *Nat. Geosci.* **12**, 685–692 (2019).
262. Kirwan, M. L. & Guntenspergen, G. R. Influence of tidal range on the stability of coastal marshland. *J. Geophys. Res. Earth Surf.* **115**, (2010).
263. Sayre, R. *et al.* Earth's coastlines. in *Wright, D. and C. Harder (eds), GIS For Science, Volume 3: Maps for Saving the Planet* vol. 3 4–27 (Esri Press, Redlands, California, 2021).
264. Easterling, D. R. *et al.* Climate Extremes: Observations, Modeling, and Impacts. *Science* **289**, 2068–2074 (2000).
265. Hijmans, R. J., Cameron, S. E., Parra, J. L., Jones, P. G. & Jarvis, A. Very high resolution interpolated climate surfaces for global land areas. *Int. J. Climatol.* **25**, 1965–1978 (2005).
266. Krauss, K. W. *et al.* Mangroves provide blue carbon ecological value at a low freshwater cost. *Sci. Rep.* **12**, 17636 (2022).
267. Title, P. O. & Bemmels, J. B. ENVIREM: an expanded set of bioclimatic and topographic variables increases flexibility and improves performance of ecological niche modeling. *Ecography* **41**, 291–307 (2018).
268. Wright, M. N. & Ziegler, A. ranger: A Fast Implementation of Random Forests for High Dimensional Data in C++ and R. *J. Stat. Softw.* **77**, (2017).
269. Kuhn, M. Building Predictive Models in R Using the caret Package. *J. Stat. Softw.* **28**, 1–26 (2008).
270. R Core Team. *R: A Language and Environment for Statistical Computing*. (R Foundation for Statistical Computing, Vienna, Austria, 2022).
271. Wenger, S. J. & Olden, J. D. Assessing transferability of ecological models: an underappreciated aspect of statistical validation. *Methods Ecol. Evol.* **3**, 260–267 (2012).
272. Roberts, D. R. *et al.* Cross-validation strategies for data with temporal, spatial, hierarchical, or phylogenetic structure. *Ecography* **40**, 913–929 (2017).
273. Ludwig, M., Moreno-Martinez, A., Hölzel, N., Pebesma, E. & Meyer, H. Assessing and improving the transferability of current global spatial prediction models. *Glob. Ecol. Biogeogr.* **32**, 356–368 (2023).
274. Milà, C., Mateu, J., Pebesma, E. & Meyer, H. Nearest neighbour distance matching Leave-One-Out Cross-Validation for map validation. *Methods Ecol. Evol.* **13**, 1304–1316 (2022).
275. Meyer, H. & Pebesma, E. Machine learning-based global maps of ecological variables and the challenge of assessing them. *Nat. Commun.* **13**, 2208 (2022).
276. Linnenbrink, J., Milà, C., Ludwig, M. & Meyer, H. kNNDM: k-fold Nearest Neighbour

Distance Matching Cross-Validation for map accuracy estimation. *EGUsphere* 1–16 (2023) doi:10.5194/egusphere-2023-1308.
277. Meyer, H., Milà, C. & Ludwig, M. *CAST: 'caret' Applications for Spatial-Temporal Models*. (2022).
278. Schratz, P., Muenchow, J., Iturritxa, E., Richter, J. & Brenning, A. Hyperparameter tuning and performance assessment of statistical and machine-learning algorithms using spatial data. *Ecol. Model.* **406**, 109–120 (2019).
279. Meyer, H., Milà, C., Ludwig, M. & Linnenbrink, J. Developer Version of the R package CAST: Caret Applications for Spatio-Temporal models. https://github.com/HannaMeyer/CAST/tree/28290c804e43251f891b7553f772a62b767b4554 (2023).
280. Pya, N. *Scam: Shape Constrained Additive Models*. (2022).
281. Mölder, F. *et al.* Sustainable data analysis with Snakemake. *F1000Res* **10**, (2021).
282. Flanders Marine Institute (VLIZ), Belgium. The union of world country boundaries and EEZ's, version 3. Marine Data Archive https://doi.org/10.14284/403 (2020).
283. Ewers Lewis, C. J. *et al.* Drivers and modelling of blue carbon stock variability in sediments of southeastern Australia. *Biogeosciences* **17**, 2041–2059 (2020).
284. Costa, M. D. de P. *et al.* Current and future carbon stocks in coastal wetlands within the Great Barrier Reef catchments. *Glob. Change Biol.* **27**, 3257–3271 (2021).
285. Hayes, M. A. *et al.* Dynamics of sediment carbon stocks across intertidal wetland habitats of Moreton Bay, Australia. *Glob. Change Biol.* **23**, 4222–4234 (2017).
286. Hughes, C. E., Kalma, J. D., Binning, P., Willgoose, G. R. & Vertzonis, M. Estimating evapotranspiration for a temperate salt marsh, Newcastle, Australia. *Hydrol. Process.* **15**, 957–975 (2001).
287. Dethier, E. N., Renshaw, C. E. & Magilligan, F. J. Rapid changes to global river suspended sediment flux by humans. *Science* **376**, 1447–1452 (2022).
